# Increased mutation rates and diversity are dominant features of *Geobacter* multi-heme cytochromes

**DOI:** 10.1101/2025.11.06.686851

**Authors:** Ruth Starwalt-Lee, Jeffrey A. Gralnick, Daniel R. Bond

## Abstract

Multi-heme cytochromes are the central catalysts of extracellular electron transfer and are uniquely abundant in the genomes of model organisms like *Geobacter sulfurreducens*. While specific functions for some multiheme cytochromes are known, the complex repertoire present in any genome makes annotation and prediction of electron transfer circuitry challenging. Here we examine patterns of conservation and rates of evolutionary change among *Geobacter* cytochromes that help explain these difficulties. Using the Ppc and OmcS cytochromes as test cases we find that sequence based methods of determining protein homology can be inadequate for distinguishing between cytochromes known to have differing functions. Importantly, using mutation rate analysis, we find that multi-heme cytochromes in *Geobacter* and *Shewanella* exhibit increased mutation rates, which may account for inaccurate homolog identification even between closely related organisms. Finally, an analysis of multi-heme cytochrome diversity reveals that each *Geobacter* genome contains a high proportion of cytochromes that are unique to that individual species, suggesting a high rate of horizontal acquisition and gene loss. These increased mutational and genetic exchange rates will need to be properly accounted for in annotation tools before we can accurately ascribe function and catalog the complex repertoire of cytochromes essential to extracellular electron transfer.

**Importance:** Dissimilatory metal reducing bacteria are found worldwide and encode diverse multi-heme cytochromes with properties suitable for applications in bioremediation, bioenergy, and bioelectronics. We find that multi-heme cytochromes involved in extracellular electron transfer show poor conservation, with significantly higher mutation rates than other elements of the proteome. This previously undescribed characteristic will limit the efficacy of standard methods of homolog annotation and database mining currently used to identify specific multi-heme cytochromes. Our findings also suggest a vast pool of undiscovered multi-heme cytochromes exists that is constantly being acquired or exchanged.

## Introduction

The discovery of extracellular electron transfer represented a significant advancement in our understanding of bacterial respiratory strategies, and uncovered a natural resource for engineering the interface between biology and electricity (*1–4*). In Gram-negative bacteria, electrons are transferred from cytoplasmic metabolism through a sequence of multi-heme cytochromes located in the cytoplasmic membrane, periplasm, outer membrane, and extracellular space, to extracellular electron acceptors such as iron or manganese oxides (*1*).

These cytochromes also allow organisms to transfer electrons to partner organisms such as methanogens, or to artificial electron acceptors like poised electrodes (*1*, *5–7*). This ability to interact with electrodes is the basis of energy-capturing microbial fuel cells, while wire-forming polymeric nanowire cytochromes are being investigated for applications in bioelectronics (*8*, *9*). Despite their environmental and technological importance, unraveling electron transfer in models such as *Geobacter sulfurreducens*, and predicting function in other uncultivated species, is slowed by the complex complement of multiheme cytochromes present in each genome.

Complicating genetic and functional analysis is the fact that many cytochromes appear to be the result of duplication events. *G. sulfurreducens* encodes several classes of cytochromes that contain up to 6 paralogs in the same genome. An instance common to *Geobacter* and its relatives is the existence of between 1 and 6 members of the Ppc family of triheme cytochromes, which are abundant periplasmic proteins needed to transfer electrons from cytoplasmic membrane proteins to cytochrome conduits in the outer membrane (*10*, *11*). Due to the fact that at least one of 5 Ppc genes (PpcA-E) are needed for extracellular electron transfer to metals by *G*. *sulfurreducens*, and they are present in nearly every Desulfuromonadales genome, they are among the most common proteins subjected to study of electron transfer function in Desulfuromonadales (*10*).

Each Ppc is differentially regulated, showing unique redox potentials and binding interactions with other cytochromes (*12–15*). Thus, correct identification of the closest relative of each well-characterized *G. sulfurreducens* Ppc homolog in other *Geobacter* genomes is essential when annotating this functional information onto other species. However, as we describe below, such identification is difficult, if not impossible.

Similarly, the conductive-wire-forming hexaheme OmcS has three other homologs in the *G. sulfurreducens* genome. Of these, only OmcS is known to form a conductive extracellular nanowire (*15*, *16*). In organisms related to *Geobacter*, identifying ‘true’ OmcS cytochromes, indicating a possible nanowire, is also challenging and prone to mistaken functional predictions. Compared to respiratory enzymes such as nitrate reductase that act as reliable markers of metabolic function during metagenomic studies, incorrect cytochrome annotation creates the risk that BLAST and Hidden Markov Model (HMM) based pipelines will generate excessive errors and limit functional prediction in metal-reducing bacteria (*17*).

This work analyzed as set of curated genomes to ask if: 1) we can reliably assign function or identify true orthologs of multi-heme cytochromes across species and genera, and 2) if poor sequence conservation is a widespread feature of cytochromes in *Geobacter*. Sequence-based homology is typically reliable for annotation and assignment of specific physiological processes across phylogenetically diverse populations. However, in our test cases using periplasmic and nanowire cytochromes of closely related organisms, this did not always hold true. As high-quality annotation is essential for data-mining genome repositories, it is critical that we understand the origins of this poor conservation before designing protein prediction tools.

Our analysis of finds evidence that increased mutation rates are a general feature of cytochromes in the model genera *Geobacter* and *Shewanella*. Further, this survey of cytochrome diversity in *Geobacter* and *Shewanella* revealed hundreds of multi-heme cytochromes that were each unique to a particular genome, lacking homologs in other strains.

The existence of a large cytochrome pangenome, evolving at rates that exceed those of standard respiratory proteins, implies an enormous sequence space is being explored by these cytochromes. This poses unique opportunities for environmental and bioelectronic applications, but will require new bioinformatic approaches to describe and understand.

## Results

### Can we distinguish ‘true’ PpcA from periplasmic triheme cytochrome homologs across Geobacter species?

Nearly every *Geobacter* genome encodes multiple triheme periplasmic cytochromes, with the largest family known by the gene designation *ppc*. In a curated set of 33 high quality *Geobacter* genomes (see Methods), genomes encode an average of 4 *ppc* genes, with a range of 1-6 (Fig. 1). In *G. sulfurreducen*s, PpcA is the best characterized and most abundant in the periplasm (*10*, *18*). When annotating a genome encoding multiple Ppcs, identifying the true PpcA ortholog is a challenge.

**Figure 1.**
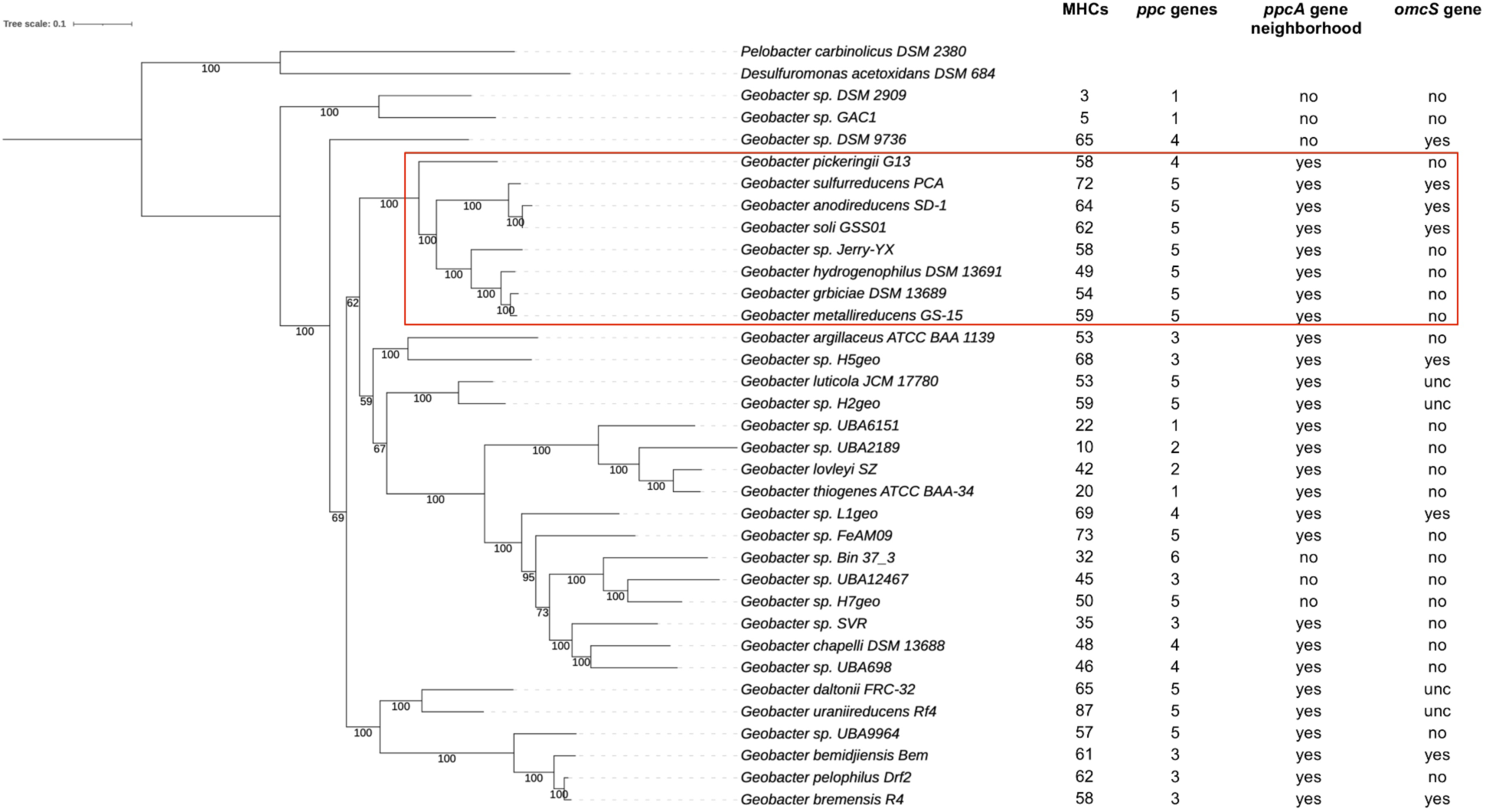
Maximum likelihood tree of *Geobacter* genomes used in this study. The box designates genomes in the ‘*G. sulfurreducens’* clade that retain the *Geobacter* genus in most current renaming schemes. Columns list the number of multi-heme cytochromes (MHCs), putative periplasmic triheme *ppc* or hexaheme *omcS* genes, and whether a conserved ‘PpcA’ gene neighborhood was found that allowed identification of putative PpcA orthologs. The tree branches are labeled with bootstrap values. Data for the *Desulfuromonas* outgroup is omitted from the analysis as only closely related strains were used for this work.

For example, two triheme Ppc cytochromes in *G. metallireducens* have higher identity to *G. sulfurreducens* PpcA than any other homolog in the *G. sulfurreducens* genome. Because these organisms are closely related, we can use conserved gene synteny around *G. sulfurreducens ppcA* to identify a likely ‘true’ ortholog, allowing the remaining highly similar gene to be designated PpcB in *G. metallireducens* based on the next best hit (Fig. 2). However, such synteny information is rarely utilized in automated bidirectional best-hit algorithms, and as PpcA is not expressed as part of a larger operon, there is little reason for synteny to be maintained. Normally, when annotation software encounters two paralogs with higher similarity to PpcA than any other protein, it is treated as a duplication, and in the case of *G. metallireducens* designates both PpcA and PpcB ‘PpcA_1’ and ‘PpcA_2’. This obscures the protein’s evolutionary history and function. The error is amplified if *G. metallireducens* acts as a reference sequence. PpcA_2 (which is actually PpcB) may be found to have a best hit to a protein, causing the new gene to be named *ppcA*. In other words, if Ppc sequences cannot be correctly identified, annotations rapidly become arbitrary.

**Figure 2.**
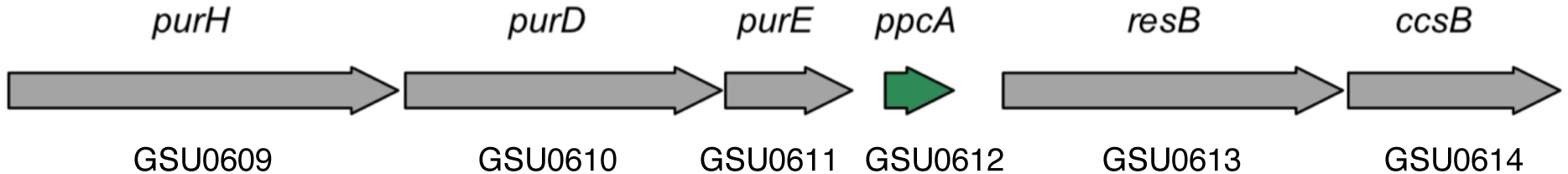
Conservation of synteny allowing identification of putative PpcA genes. The conserved *G. sulfurreducens ppcA* gene neighborhood.

To see how Ppc homologs challenge annotation, we tested the ability of standard bioinformatic approaches to identify PpcAs using both a genome annotation pipeline and a homolog clustering pipeline. First, Prokka was used to re-annotate a selection of high-quality *Geobacter* genomes, using our manually annotated *G. sulfurreducens* PCA genome as the reference (*19*). In this workflow, Prokka identified all putative coding sequences, then annotated them based on their bidirectional best BLAST hit to the reference *G. sulfurreducens* genome.

Of 121 *ppc* annotations found across 33 genomes, 64 were annotated as *ppcA*. In 21 cases, between 2 and 4 genes were designated ‘*ppcA*’, while only nine genomes contained a single cytochrome annotated as *ppcA*. As expected, when we manually inspected genomes, most *ppcA ‘*duplicates’ did not share synteny with *G. sulfurreducens* PpcA, indicating they were not true orthologs (Fig. 2). The sequences of the Ppc cytochromes were in similarity ranges (pairwise similarity ∼ 35-100% with BLOSUM62) that prevented clear bidirectional best hits to be resolved, causing them to be grouped together and called the same protein.

These annotated *Geobacter* genomes were then analyzed with a bioinformatic pipeline (get_homologues), which creates homolog clusters from all-vs-all genome BLAST data using the Ortho Markov CLuster (OMCL) algorithm (*20*, *21*). OMCL is not limited to pairwise comparisons, and because it considers an overall similarity network, it tends to be more sensitive in identifying orthologous genes in the presence of paralogs. This clustering is also better suited to variations in evolutionary distances between genes, making it more effective across a range of species divergence (*21*).

Using the OMCL algorithm, the majority of triheme cytochromes (58 of 121) still grouped into one large cluster with the *G. sulfurreducens* PpcA, while the remainder separated into 16 homolog clusters. Of the 58 cytochromes clustered by OMCL with *G. sulfurreducens* PpcA, 45 had also been designated as PpcA by bidirectional best-hit annotation. The remaining proteins annotated by best-hit as PpcA were scattered in other clusters by OCML, with no clear pattern for PpcB-E.

We then asked if signature residues or motifs could allow more sensitive comparisons. For example, sequences from the 58-member OMCL PpcA cluster were aligned and used to construct a Hidden Markov Model (HMM). Using the pipeline FeGenie, which utilizes user-supplied HMM models and bitscore thresholds, we queried the original set of 33 genomes, accepting any hit over an intentionally low bitscore of 100 (*22*). This ‘OMCL PpcA-cluster’ HMM retrieved 75 sequences (Fig. 3A). While 85% of the sequences used to construct the HMM scored higher than other sequences, there was no obvious bitscore cutoff that could distinguish members of the original PpcA pool from non-members. In fact, sequences clustering with *G. sulfurreducens* PpcB or PpcE scored very well using the PpcA HMM in the FeGenie pipeline, as did members of variant triheme proteins lacking any *G. sulfurreducens* homologs. In other words, an HMM constructed out of an aligned cytochrome collection, that was supposed to only identify that subset, instead scored about 60% of all *Geobacter* triheme cytochromes well, and would have named all paralogs within the *G. sulfurreducens* genome PpcA. This did not appear to relate to the physiological function or evolutionary history of these proteins.

**Figure 3.**
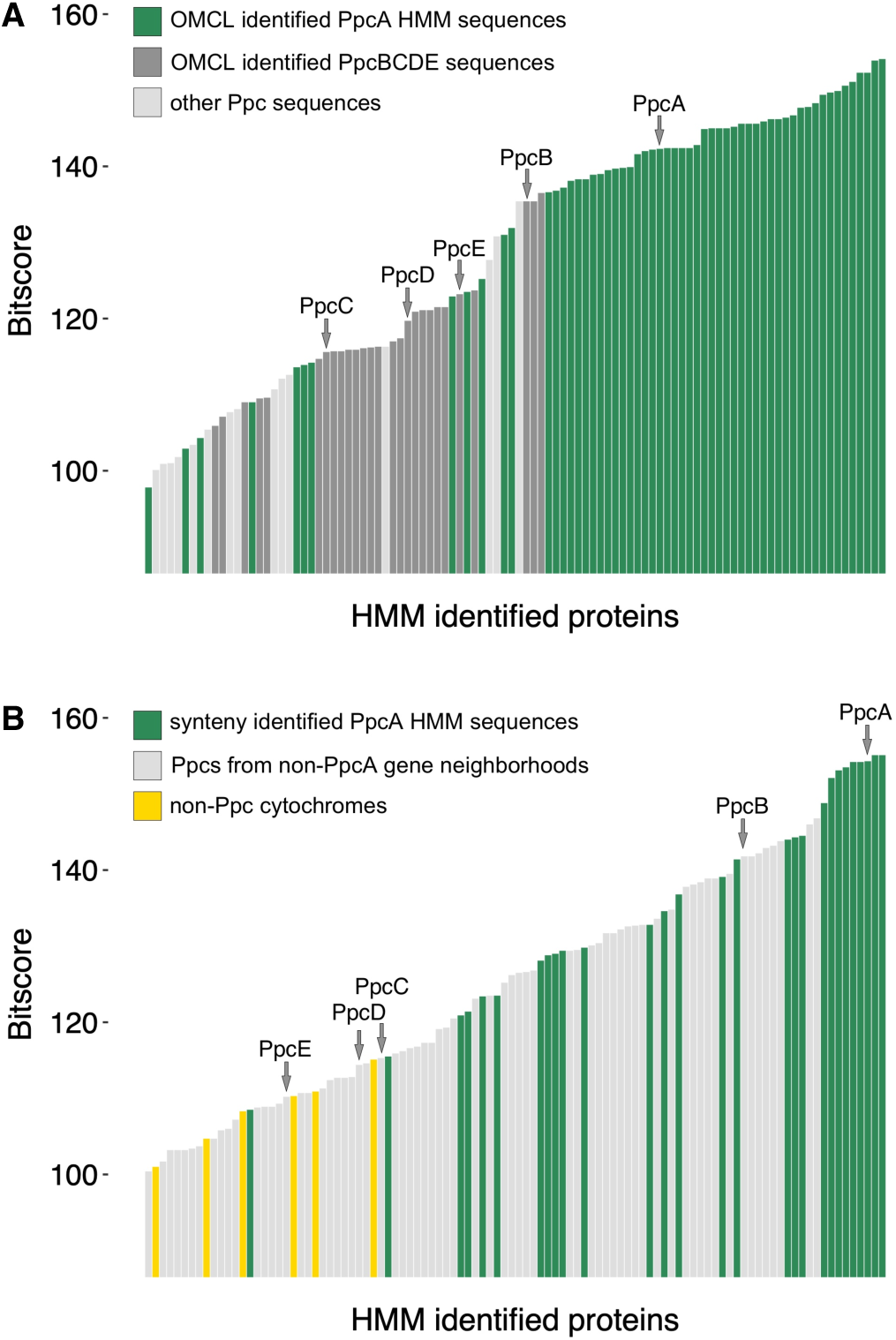
Neither homology nor synteny based HMMs of PpcA can distinguish PpcA from other Ppc paralogs. A) Results from the homology-based HMM, using PpcA sequences clustered by OCML with *G. sulfurreducens* PpcA. Sequences designated PpcBCDE were clustered by OMCL with *G. sulfurreducens* PpcB, C, D, or E. B) Results from the synteny-based PpcA HMM, using manually curated PpcA sequences. Ppc representatives from *G. sulfurreducens* PCA are designated by arrows.

As we previously noted, there is conserved synteny around some *ppc* genes, especially in strains closely related to *G. sulfurreducens*. Based on this synteny, we tried a manual approach to creating a ‘true’ PpcA collection, that could train an HMM (Fig. 3B). We hypothesized that synteny-confirmed sequences would provide a set of authentic PpcA sequences with shared ancestry, with the best chance of identifying signature residues. Twenty-eight sequences were identified for this synteny-verified PpcA HMM.

Using the FeGenie pipeline, the HMM built from the careful synteny-based PpcA collection retrieved an even wider, and even less accurate, collection of cytochromes. This approach, designed to only identify *ppcA* genes, scored *G. sulfurreducens* PpcB higher than 16 of the ‘verified by synteny’ PpcA sequences from other genomes. Non-*ppcA* homologs within the *G. sulfurreducens* genome (PpcB-E) also scored well (Fig. 3B). This HMM retrieved multiheme cytochromes that were not even related to the Ppc family. Once again, there was no obvious bitscore cutoff that could be programmed into FeGenie to allow discrimination of PpcA from homologs separated into distinct clusters by OMCL, or which had been annotated as non-PpcA cytochromes during annotation with Prokka.

In these examples, evidence suggests these triheme cytochrome sequences do not maintain enough global or local sequence conservation for standard tools to provide reliable annotation outside of belonging to a triheme family. Whether it was standard BLAST best-hit, more robust ortholog cluster algorithm, or manually curated HMM using authentic homologs identified by gene synteny, each approach produced different and ambiguous results. Any of these methods, would be useful for identifying triheme cytochromes mostly belonging to the Ppc family, but not for assigning orthology or transferring a true name like PpcA or PpcB.

One explanation for the failure of sequence based methods in this example is that the Ppc family of cytochromes are “peptide-minimized” (*23*). These ∼9-10 kDa/3 heme proteins are near the lower limit of how much peptide is needed to fold around and coordinate three hemes. Most of the evolutionary pressure may be focused on maintaining the heme fold, with few residues involved in physiologically relevant interactions. This is not just a feature of small cytochromes, as many larger multiheme cytochromes, have a similar peptide-minimized ratio; such as the ∼70 kDa/26 heme GSU2495, 32 kDa/12 heme ExtA, and 25 kDa/8 heme OmaB. However, larger cytochromes, especially those with a higher amino acid:heme ratio are more likely to contain more useful information.

### Can we identify OmcS nanowires?

The *Geobacter* OmcS cytochrome that polymerizes into an extracellular nanowire poses a problem similar to the Ppc cytochromes. In *G. sulfurreducens* there are three hexaheme cytochromes related to OmcS (GSU2504). Two of these, OmcT (GSU2503), and an “OmcS-like” unnamed cytochrome (GSU2501), are encoded within an operon with *omcS*, while *omcJ* (GSU0701) is elsewhere. To date, the functions of *G. sulfurreducens* OmcS homologs are unknown, and only OmcS is known to form a wire, making it important to differentiate OmcS from its relatives (*16*). To capture all possible homologs, we collected all hexa-heme protein sequences within 10% of the *G. sulfurreducens* OmcS length from our 33-genome set. Unlike the PpcA example, where every genome contained multiple homologs, only 12 genomes had any cytochromes related to OmcS. The presence or absence of an *omcS*-like gene in a genome did not correlate with species phylogeny, providing evidence that this group is subject to frequent loss and gain.

A gene phylogeny of this *omcS* collection showed evidence of multiple independent duplication events within the family (Fig. 4). For example, *G. sulfurreducens omcS*, *omcT* and *omcJ* appear to have arisen from an ancestral hexaheme cytochrome. The phylogeny of these *omcS*, *omcT* and *omcJ* copies is similar to a species tree of the strains containing these genes (Fig. 4, Clades 1 and 2). However, instead of containing *omcT* and *omcJ*, *G. bemidjiensis* contains three copies of *omcS* that are more closely related to each other than to any other homolog, suggesting a new series of duplication events happened more recently. It is not clear whether each new *omcS* still encodes a nanowire subunit (Fig. 4, Clade 2).

**Figure 4.**
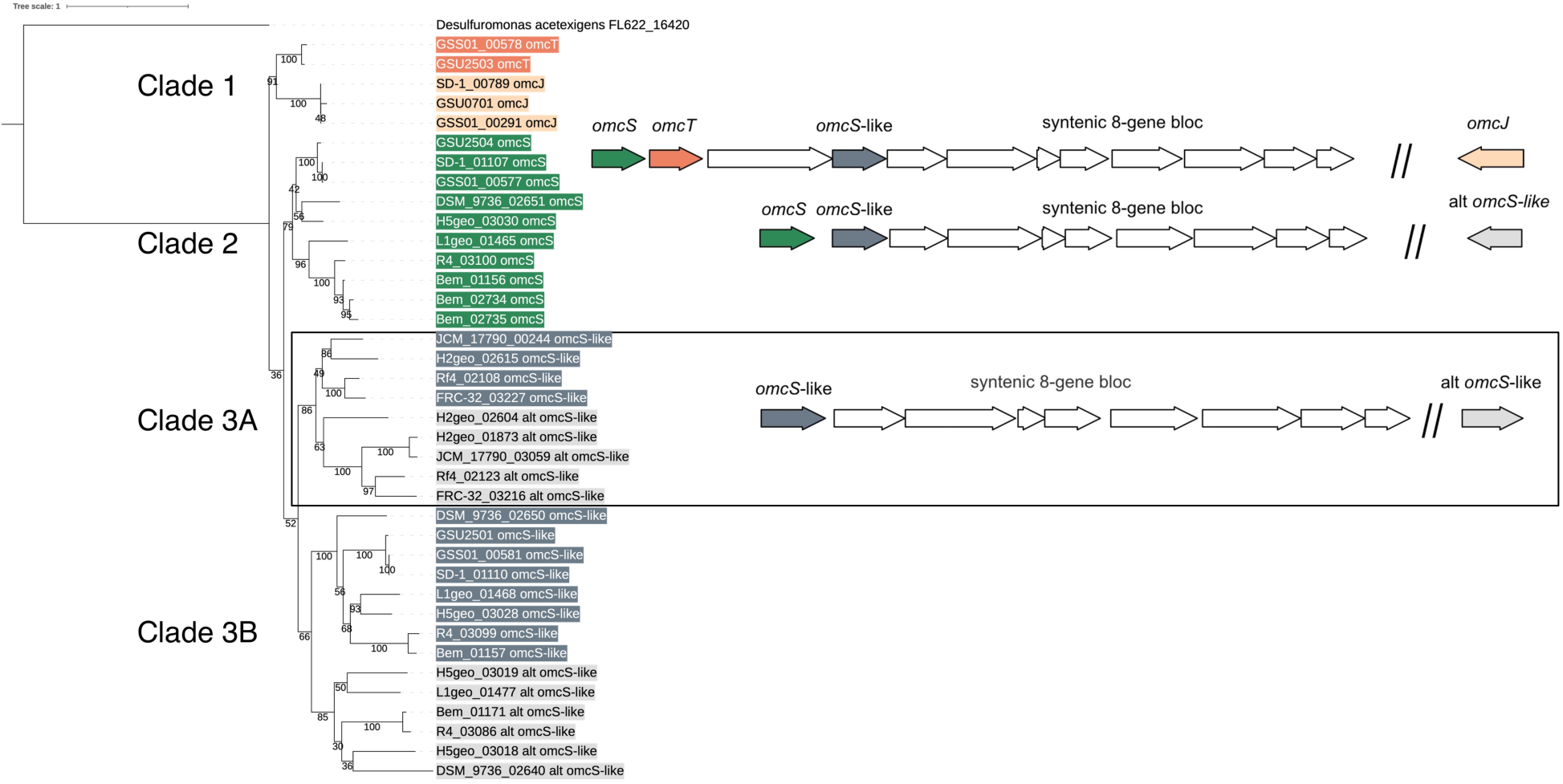
Phylogeny and gene synteny of *Geobacter* genes homologous to *G. sulfurreducens omcS*. Gene neighborhoods are color-coded to correspond to the gene phylogeny. The top gene neighborhood is found in *G. sulfurreducens*, *G. anodireducens*, and *G. soli*, with genes from Clades 1 and 2. *G. soli* and *G. anodireducens* have identical *omcT* sequences, so *G. anodireducens omcT* was removed from analysis. The middle gene neighborhood is found in *G. bemidjiensis*, *G. bremensis*, *G.sp.* DSM_9736, *G. sp.* H5geo, and *G. sp.* L1geo, with genes from Clades 2 and 3B. The boxed gene neighborhood is found in *G. daltonii*, *G. luticola*, *G. sp.* H2geo, and *G. uraniireducens*, with genes exclusively from Clade 3A.

Cytochromes encoded by the “*omcS*-like” hexaheme GSU2501 that usually occurs near *omcS* do not cluster with *G. sulfurreducens omcS*, suggesting these diverged longer ago. Interestingly, all genomes with this “*omcS*-like” GSU2501 homolog have an eight gene block downstream (*G. sulfurreducens* GSU2500-GSU2493) recently linked to OmcS nanowire protein maturation and secretion (46). While not all 12 genomes containing the eight-gene ‘nanowire secretion’ cluster encode a *G. sulfurreducens omcS*, all encode the GSU2501 hexaheme cytochrome. This suggests GSU2501 could be ancestral to OmcS, and forms an uncharacterized nanowire, or GSU2501 is part of a cluster that was later adopted for nanowire production when OmcS arose. Overall, hexaheme OmcS and its homologs are encoded in diverse patterns (Fig. 4): 1) an operon containing *omcST* and the GSU2501 *omcS*-like gene (sequences in Clades 1 & 2) *omcS* and a GSU2501 *omcS*-like gene (sequences in Clades 2, 3B), or 3) only a GSU2501*omcS*-like gene upstream of the eight gene block (sequences in Clade 3A). Further, additional duplications remain unique to individual species, such as the hexaheme cytochromes elsewhere on the *G. bemidjiensis* genome.

Like our efforts with PpcA, we tested whether homology-based or gene synteny-based alignments could identify ‘authentic’ OmcS homologs, or create HMMs capable of distinguishing these different hexaheme cytochrome families. For the homology-based identification, a small get_homologs cluster containing a subset closely related to *G*.

*sulfurreducens* OmcS was used to create an HMM. Unlike the results for the triheme cytochromes, the small number of input sequences used to create the HMM were correctly retrieved with the highest scores and were more distinct from OmcT, OmcJ, and other OmcS-like sequences (Fig. 5A).

**Figure 5.**
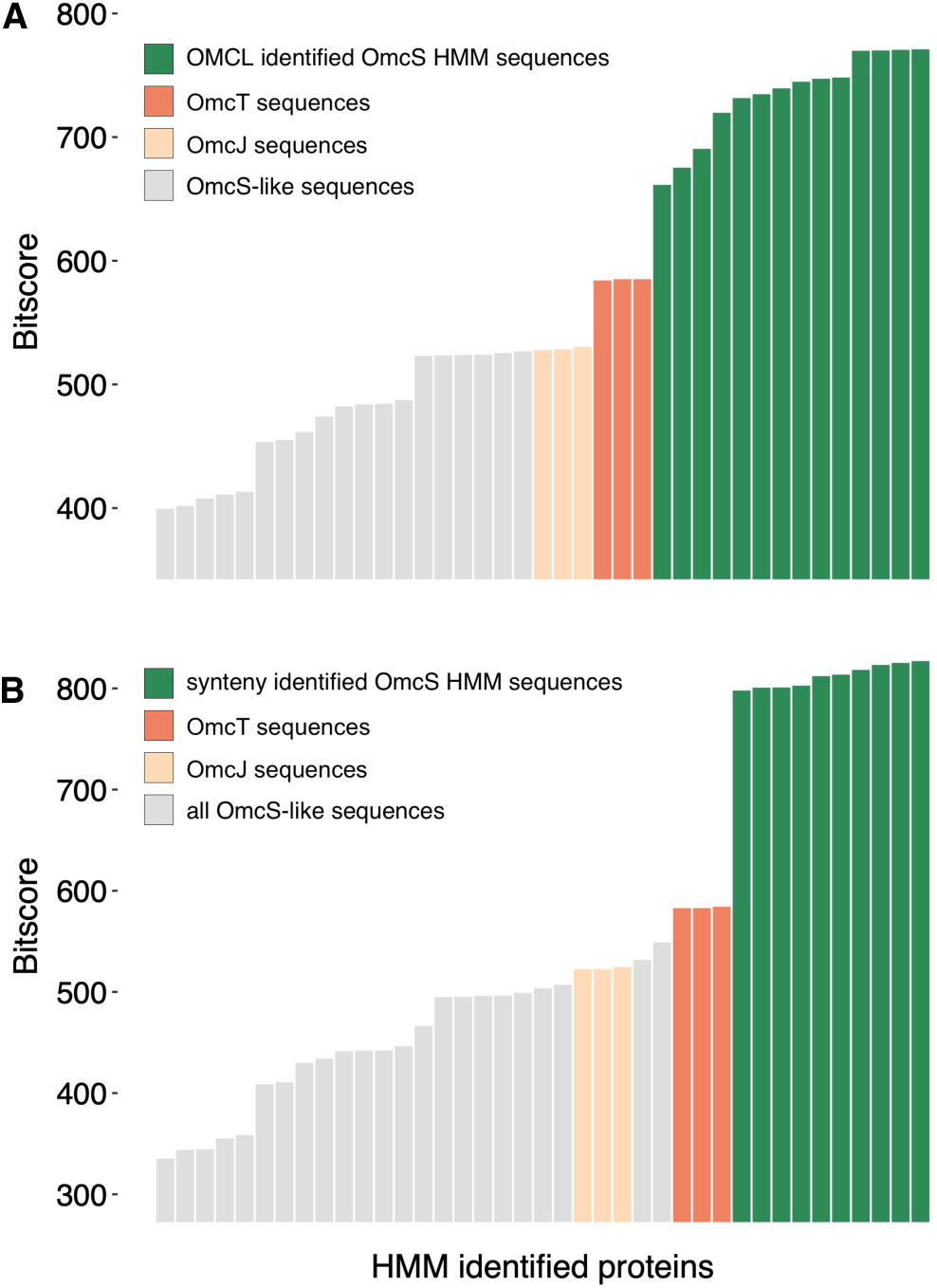
Homology and synteny based OmcS HMMs are more successful at distinguishing OmcS from related 6-heme homologs. A) Results from the homology based OmcS HMM built from OMCL clustered OmcS. B) Results from the manually curated synteny based OmcS HMM.

When OmcS sequences were carefully selected using synteny, the resulting HMM-based identification was even more restrictive, notably clustering OmcT or OmcJ with the GSU2501 OmcS-like sequences (Fig. 5A). Together, these results suggest that that, unlike with Ppcs, there may be signatures within *very close relatives* sufficient to distinguish OmcS from OmcT and OmcJ or other hexaheme cytochromes. We stress that such HMMs are likely only useful when searching within a genus, and would lack the resolution needed to detect new OmcS nanowires in other genera.

OmcS and its homologs have higher kDa/heme ratios, which indicates there may be residues at active protein-protein interaction sites in addition to their heme-binding regions. Lacking clear information about which subunits form nanowires or catalyze other functions, sequence selection for HMM creation remains difficult. In a case like *G. bemidjiensis*, recent duplications created homologs that could not be distinguished by our methods without new functional information. The fact that only a few genomes contain OmcS, while others have multiple duplications, and occasional genomes have hexaheme cytochromes of unknown function again highlights how these proteins appear to be rapidly changing.

### Is there evidence for greater mutation rates in Geobacter cytochromes?

The continuum of low bitscores from our alignments and HMMs suggested that the sequences of multi-heme cytochromes changed faster than the average protein. To test this hypothesis, we performed mutation rate analysis on curated cytochrome homologs from our *Geobacter* genome set, as well as a set of *Shewanella* genomes.

In this analysis, the ratio of non-synonymous nucleotide changes (dN) to synonymous nucleotide changes (dS) acts as a measure of mutation rate over time. Since synonymous mutations are less affected by selective pressures than non-synonymous mutations, they can act like a clock, normalizing comparisons between genes separated by different amounts of evolutionary time (*24*).

As both types of mutations should occur during DNA replication at the same rate, a dN/dS ratio of 1 would occur with no selection. If the protein sequence undergoes selection to retain the same residues, non-synonymous mutations should not remain fixed in the population, and show a slower rate of amnio acid changes relative to the baseline synonymous mutation rate. A selection to remain the same manifests as a low dN/dS ratio. Selection for the protein to adapt, or a lack of selection allowing drift, would cause the dN/dS to increase. This metric has the benefit of standardizing comparisons between closely related and distant species, assuming their baseline DNA mutation rate is the same.

In these analyses, dN and dS values were obtained by comparison of a reference gene in *G. sulfurreducens* PCA to its verified homolog in another *Geobacter* genome. *Shewanella oneidensis* MR-1 was used as the reference for *Shewanella*. Briefly, 33 *Geobacter* species and 39 *Shewanella* species were analyzed to obtain homolog clusters (Table S1). Because of problems described earlier with highly duplicated cytochromes such as Ppc or OmcS cytochromes, we eliminated genes that contained unresolvable paralogs in each genome.

Clusters containing a reference PCA or MR-1 sequence were manually curated as follows: 1) the homolog sequence length had to be within 20% of the reference protein, 2) the number of heme binding motifs had to be ± 2 of the reference, and 3) if a cluster contained two or more homologs from the same genome, the correct cytochrome homolog had to be manually verified, or else the sequences were removed from further analysis.

Pangenome matrices derived from all *Geobacter* or *Shewanella* genomes also were used to identify conserved genes present in all genomes, designated as “core” genes. Protein sequences aligned with Clustal Omega were then used to codon format corresponding nucleotide sequences, so that mutation rates for each homolog could be obtained by pairwise comparisons of each sequence using the KaKs Calculator (*25*, *26*).

The results of these mutation rate analyses show that, for the majority of analyzed *Geobacter* genomes (25/32), mutation rates are significantly higher for cytochrome genes than core protein-encoding genes (Fig. 6A, Fig. S1). As many cytochromes with high mutation rates (high dN/dS ratios) had to be removed from our final analysis due to the potential for erroneously comparing paralogous sequences, the magnitude of this mutation rate is likely underestimated.

**Figure 6.**
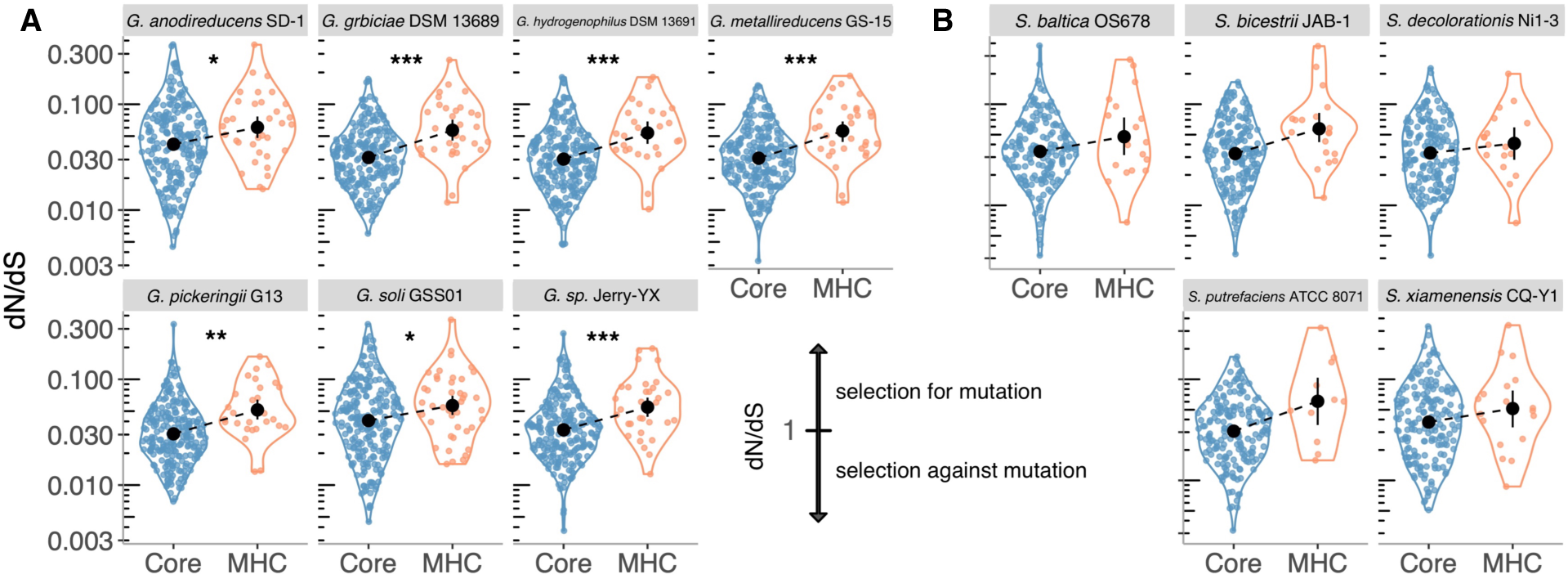
Comparison of core vs. multiheme cytochrome mutation rates in *Geobacter* and *Shewanella*. Higher values represent faster rates of non-synonymous amino acid replacements. A) Core vs. multiheme cytochrome mutation rates from genomes in the *Geobacter sulfurreducens* clade. B) Core vs multiheme cytochrome mutation rates from genomes in the *Shewanella oneidensis* clade. Significant differences are designated by asterisks: * < 0.05, ** < 0.001, *** < 0.0001, ns – not significant

The results for *Shewanella* also showed 16/38 genomes had statistically significant differences (Fig. 6B, Fig. S2). As *Shewanella* do not have as many cytochromes as *Geobacter,* insufficient data may contribute to the lack of statistically significant differences between core and cytochrome genes in *Shewanella*. Notably, five *Geobacter* genomes that did not have statistically significant differences between core and multi-heme genes also were strains with the fewest multiheme cytochromes (<22, Fig. S1).

While direct comparison of *Geobacter* dN/dS rates to *Shewanella* dN/dS rates is not possible as they do not have any cytochromes in common, mutation rates were highest in *Geobacter* cytochromes. Although many *Geobacter* genomes have since been assigned to other Geobacteraceae genera, all genes in this analysis were standardized to the synonymous mutation rate. This means the enhanced mutation rates in our data set are not due to larger phylogenetic distances between *G. sulfurreducens* and genomes reassigned to new genera, but are inherent characteristics of these proteins.

While *Geobacter* cytochromes show greater mutation rates compared to other conserved proteins, we observed that mutation rates for cytochromes were not uniformly high. We hypothesized that the localization or function of cytochromes within the electron transport chain could be correlated with mutation rate, and reanalyzed dN/dS data to test this hypothesis. For *Geobacter*, in order to include carefully annotated periplasmic *ppc* genes in this analysis, we used data only from the ‘*G. sulfurreducens* clade’ or genus, to use only closely related strains, where orthologs could be verified (Fig. 1). Analysis of dN/dS ratios showed three distinct groups in both the *G. sulfurreducens* and *Shewanella data*: 1) “low” mutation rates, typically exhibited by inner membrane cytochrome genes, 2) “medium” mutation rates, exhibited by non-*ppc*/*fccA*/*cctA* periplasmic cytochrome genes, and 3) “high” mutation rates, exhibited by *ppc*/*fccA*/*cctA*, outer membrane, and extracellular cytochrome genes (Fig. 7).

**Figure 7.**
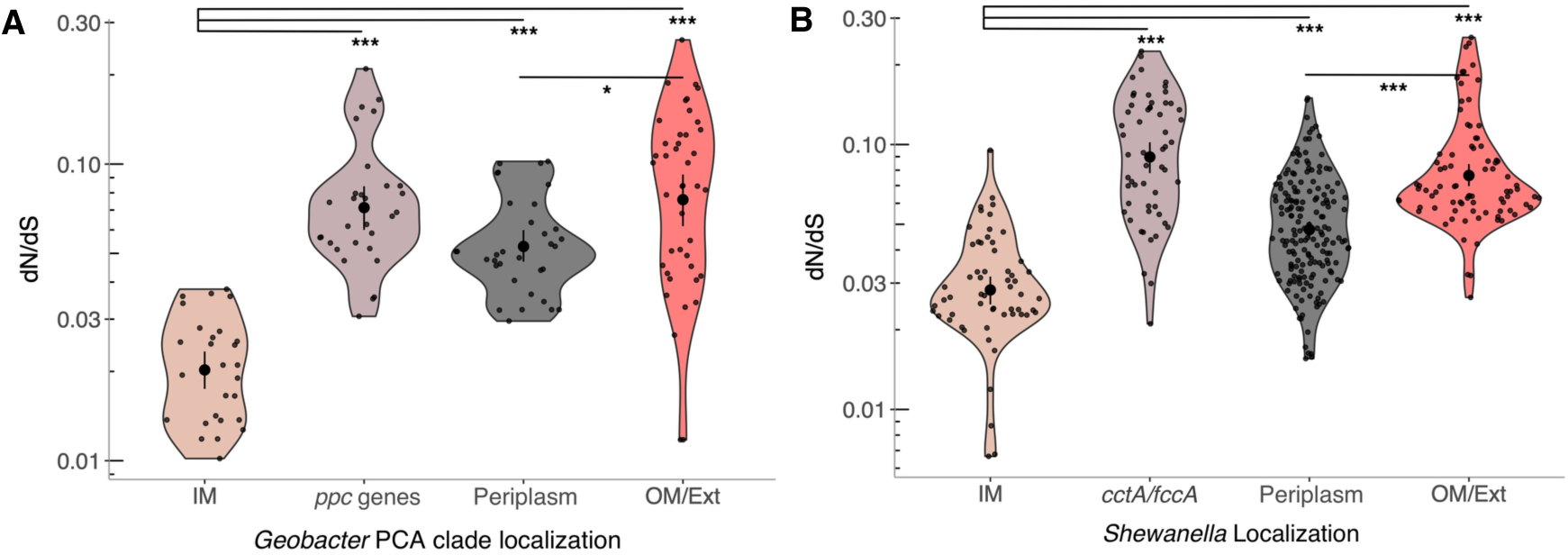
Comparison of multiheme cytochrome mutation rates in different cell locations in *Geobacter* and *Shewanella*. A) Multiheme cytochrome mutation rates for orthologs in genomes within the *Geobacter sulfurreducens* clade. IM: *imcH, cbcL*, *cbcBA*, *ppc*: *ppcABCDE*, periplasm: *nrfA*, *nrfH*, *extA*, *omaC*, *extK*, OM/ext: *omcE*, *omcS*, *omcZ*, *pgcA*, *extD*, *extC*, *omcC*. B) Multiheme cytochrome mutation rates for *Shewanella* genomes. IM: *cymA*, *torC*, periplasm: *nrfA*, *otr*, *sirA*, *mtrA*, *mtrD*, *dmsE*, OM: *mtrC*, *mtrF*, *omcA*

Of interest is the fact that genes encoding ‘periplasmic’ cytochromes that are part of larger porin-cytochrome complexes, as well as periplasmic cytochromes with strictly enzymatic functions had lower dN/dS ratios compared to *ppc*, *cctA*, and *fccA* genes. PpcA, CctA, and FccA are presumed to be free-floating in the periplasm and, with the exception of FccA, which is also a fumarate reductase, their functions are limited to electron transfer (*27*). Overall, our results indicate that the location or function of cytochromes can influence their evolutionary rate, making identification of some more reliable than others.

### The expanding cytochrome pangenome

During our analyses, we noted that *Geobacter* species have large cytochrome repertoires that undergo gain, loss, duplication, and greater mutation rates, so we asked whether a consensus repertoire can even be defined. For the 33 *Geobacter* genomes analyzed (Table S1), there was only one cytochrome with sequences in all genomes, the PpcA sequence, with the caveat that absolute identification of PpcA homologs remains questionable. Many of the analyzed genomes have been reassigned into new genera, meaning there are no cytochromes conserved across the roughly 3 genera analyzed in this work.

Across the 33 *Geobacter* genomes, a large number of multiheme cytochromes were found in only one genome (53% of OCML clusters). Cytochromes found in only 2-9 genomes represented another large percentage (31%). Only 16% of cytochromes identified contained homologs in more than 10 genomes (Fig. 8A). This suggested that not only are cytochromes evolving at rapid rates, but that each *Geobacter* genome contains a significant number of novel cytochrome sequences.

**Figure 8.**
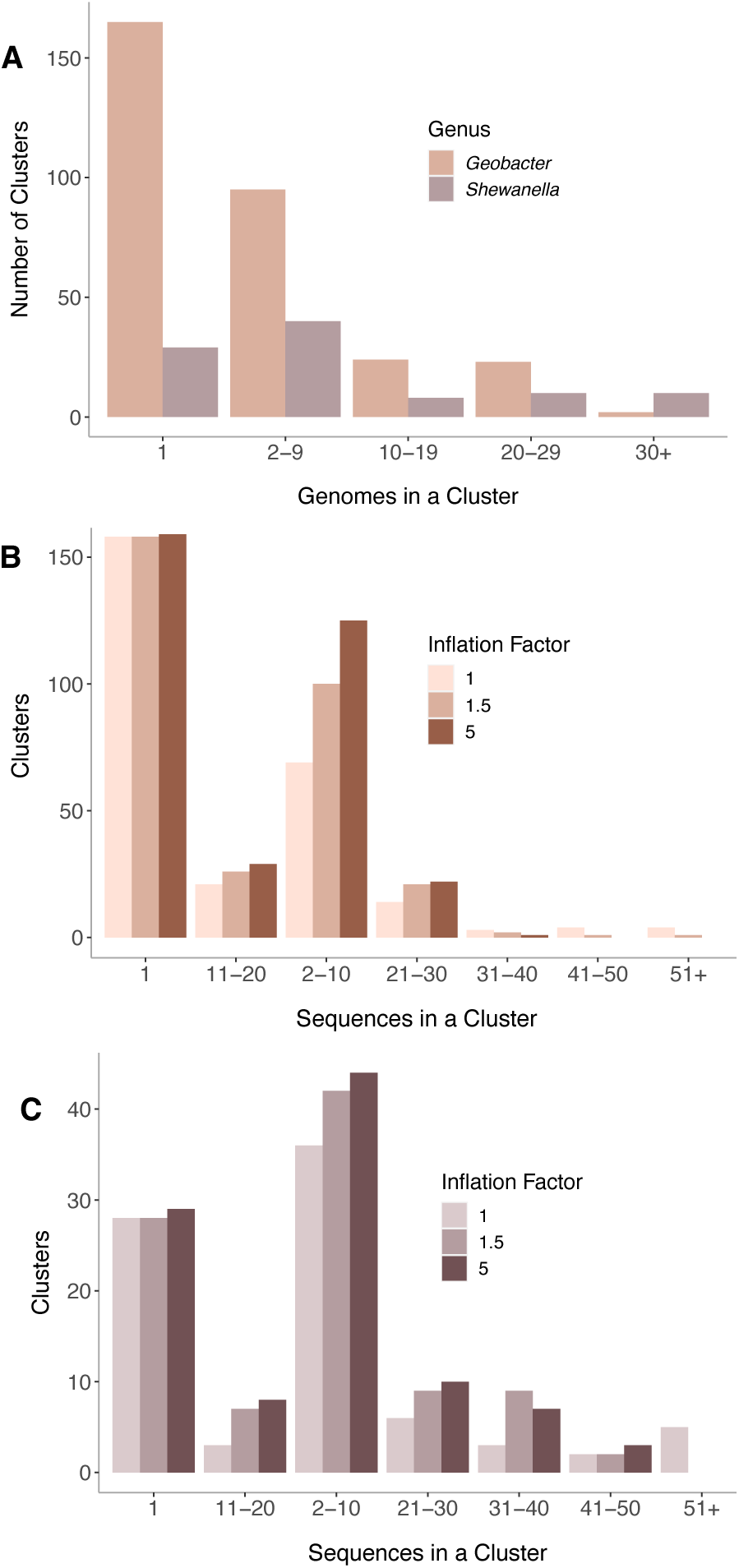
OMCL-based clusters of multi-heme cytochrome sequences are dominated by clusters with very few members. A) *Geobacter* cytochrome homolog clusters binned by the number of genomes containing a representative from a cluster. Over 150 cytochromes were found that lacked a homolog in any other *Geobacter* genome. B) Reducing homolog-calling stringency does not affect the number of *Geobacter* homolog clusters containing only one member. C) Reducing homolog-calling stringency does not affect the number of *Shewanella* homolog clusters containing only one member. Legend: F1 – lowest stringency, F1.5 – default stringency, F5 – highest stringency.

As this result could be an artifact of the stringency of our homolog calling, we repeated the analyses using the highest and lowest stringency settings.

Changing the stringency had a negligible effect on the number of unique multiheme cytochromes identified. As expected, at lower stringency, more clusters of paralogs collapsed together, grouping most PpcA-like or OmcS-like sequences into a single group (Fig. 8B). This supported the conclusion that while genomes can have many cytochromes due to duplication, the high number of novel multi-heme cytochromes, in particular the unique ‘singletons’ found in each genome, is not an artefact of the clustering algorithm.

The same analyses in *Shewanella* found that the proportion of cytochromes shared across the genus was greater than in *Geobacter* (Fig. 8A,C). Twenty percent of homolog clusters were found in more than half of the genomes analyzed (Fig. 8A). This could partially be due to the analyzed *Shewanella* genomes being more closely related. However, we found that clusters containing only one sequence constituted 30% of the total pool, as compared to 53% for *Geobacter* (Fig. 8C). The lower proportion of singleton clusters, in combination with the lower absolute numbers of cytochromes, shows that the *Shewanella* multi-heme cytochrome pool is less diverse than in *Geobacter* and related genera.

This raised the question of what level of sampling of *Geobacter* and *Shewanella* genomes would saturate the pool of cytochromes in their respective genera. For this analysis, we calculated the phylogenetic distance of each genome from the reference genome (*G. sulfurreducens* PCA or *S. oneidensis* MR-1) using a whole-genome maximum likelihood approach (*28*). We then conducted clustering analyses by adding genomes one by one, in order of least to greatest phylogenetic distance from the reference. If sampling is approaching saturation, adding a new genome should not add any unique cytochromes. The number of clusters discovered with each new genome should plateau.

For *Geobacter*, the number of new clusters continued to increase with the addition of each new genome, while clusters in *Shewanella* were beginning to taper (Fig. 9A). Given that half of the cytochromes in our *Geobacter* analysis were unique, lacking any homologs in other genomes, it is not surprising that we did not see saturation. When we removed these unique ‘singleton’ sequences, the number of cytochromes shared by two or more genomes did begin to reach saturation (Fig. 9B). These results suggest that there are multiple pools of cytochromes with different evolutionary trajectories. Many are found in 30-50% of genomes. These are constantly being lost, gained, or duplicated, and are likely frequently exchanged between organisms. But the reservoir, and function, of the much larger collection of cytochromes being infrequently horizontally acquired and lost remains unknown.

**Figure 9.**
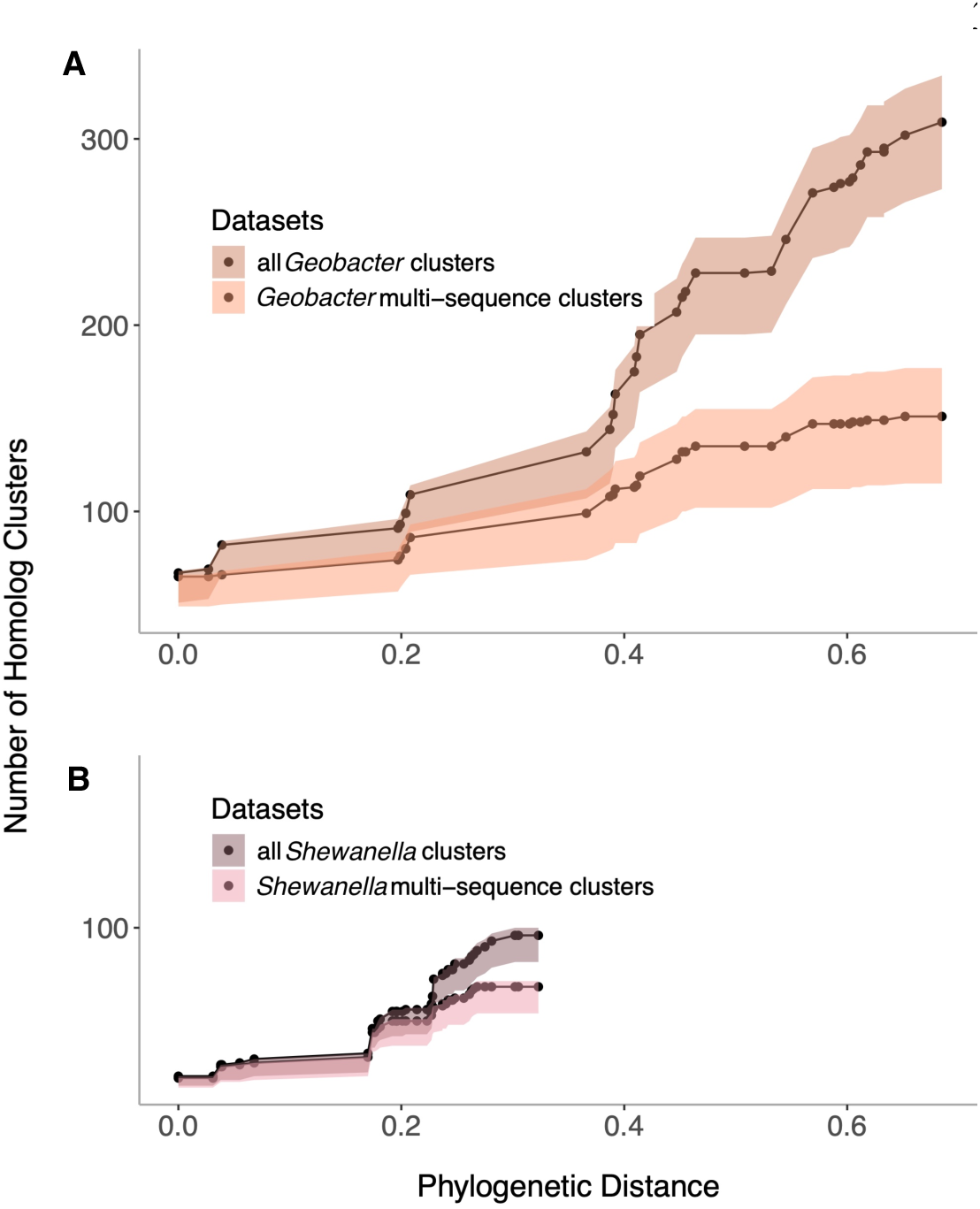
The number of unique multiheme homolog clusters continues to increase with addition of new genomes. Analysis showing the number of total cytochrome clusters as new *Geobacter* (A) and *Shewanella* (B) genomes are added. Counting only clusters with sequences found in two or more genomes shows how the increase in new families is driven by the presence of novel multi-heme cytochromes lacking any homolog in any genome.

## Discussion

The reductases of most anaerobic energy generation strategies are well conserved and easy to identify in genomes. Metagenomes can quickly be annotated to predict if organisms participate in cycling of acceptors such as nitrate, nitrite, sulfate, polysulfide, arsenate, selenate, chlorinated compounds, DMSO, TMAO, and carbon dioxide. Since the first genomes of dissimilatory metal reducing bacteria were sequenced, it has been clear that something about the proteins central to their metabolism is different.

Many key cytochromes essential to extracellular electron transfer are now known in model strains. When annotating future genomes and transferring this growing body of functional information, the assumption is made that these sequences and their functions are conserved.

However, the presence of abundant paralogs, lack of cytochromes conserved between species, and evidence for increased mutation rates raises questions about whether information from one strain should be mapped in automated bioinformatic pipelines (*1*, *10*). For example, in this study found that in families like PpcABCDE and OmcSTJ, the identification or assignment of function across broader *Geobacter* genera faces significant hurdles and should not be done by bidirectional best hit or HMM-based pipelines.

While duplications and paralogs are known issues in metal-reducing bacteria, the mutation rates of these proteins remained largely unexplored. A high mutation rate was reported within multi-heme cytochrome sequences when *G. sulfurreducens* PCA was compared to the *G. sulfurreducens* variant KN400. While these strains had 96% of coding sequences in common, and 72% of single-nucleotide differences were synonymous mutations, 25 orthologs had notably high rates of non-synonymous changes. This pool was significantly enriched in *c*-type cytochromes (*29*). In our extended analysis, this increased cytochrome mutation rate was a consistent trait across known *Geobacter*.

While multi-heme cytochromes had higher mutation rates in both *Geobacter* and *Shewanella* genomes, we found this mutation rate to be correlated with cytochrome location. Inner membrane-encoding genes have the lowest mutation rates, especially in their membrane domains, which may be due to additional sequence constraints imposed by bioenergetic coupling, reacting with the quinone pool, and interacting with periplasmic partners. These quinone oxidoreductases, such as ImcH, CbcL, CbcBA, and CymA remain among best markers for identifying putative extracellular electron transfer in metagenomic studies, due to their slower mutation rates and lower frequency of loss from genomes.

In contrast, genes from outer surface cytochromes demonstrate the highest mutation rates. A basis for these higher rates was suggested by the structures of cytochrome nanowires, where there is a high degree of structural alignment between hemes. For example, the four hemes of OmcE and four hemes in OmcS are superimposable, even though these nanowires lack any sequence similarity (*15*, *30*). Both proteins lack substantial amounts of secondary structure, allowing mutations, insertions, and deletions to easily occur in loops and coils, so long as they do not perturb the alignment and coordination of hemes (*15*, *30*). If the primary selective pressure on a cytochrome is to maintain the same heme coordination, much more variation is possible. In the case of surface exposed cytochromes, recent data showing that the *Shewanella* OmcA cytochrome is a phage target indicates pressure for phage resistance may create additional selection for amino acid changes in extracellular cytochromes (*31*).

Unlike enzymes with highly evolved active sites, where specific residues are essential, multi-heme cytochromes may be able to perform the same job with vastly different amino acids, providing they: 1) have the correct heme coordination, 2) have the correct redox potential, and 3) have a surface charge that lets them get close enough to their partner for electron transfer to occur. In the case of *G. sulfurreducens*, successfully transferring electrons across the periplasm only requires sufficient expression of any one of the five Ppc homologs, and this function can even be performed by a cytochrome typically secreted to the outer surface (*10*). The fact that increased cytochrome mutation rates are also seen in *Shewanella* suggests that such selective pressures, and promiscuous behavior, is not unique to *Geobacter*. In this case, periplasmic cytochromes will provide unreliable markers for extracellular electron transfer and be consistently misannotated.

Unlike most reductases that form the basis for most anaerobic respirations, whose presence or absence can signal an ability, the cytochromes of *Geobacter* represent an opposite situation. The number of cytochromes per genome is high, but the a low percentage of cytochromes shared between any two strains means there are few to use as markers. Every sequenced genome reveals new cytochromes not present in any other strain, raising the question of where these unique proteins are coming from? Are these unique cytochromes the result of such rapid evolution we can no longer recognize similarity with their ancestors, or is there an abundance of unique proteins being exchanged via horizontal gene transfer from species yet to be isolated?

A similar categorization dilemma, and abundant unclassifiable cytochrome problem was raised in a recent survey of a much larger collection of *Geobacter-*related genomes (45). In this survey of all available *Desulfuromonadia*, the inability to discern paralogs required treating all hexaheme OmcS, OmcT, and OmcJ-like cytochromes as similar. This allowed finding whether a genome did, or did not, have between 0 and 5 *omcS*-like genes, but prevented deduction of whether nanowire-like functions were present. In addition, even when using a MMSeqs-based approach less stringent that what was applied in our work, clustering proteins with 25% sequence identify over 70% of shared length with known cytochromes, nearly half of 4,716 cytochromes had no known homologs in other databases.

While clustering algorithms parse a large percentage of cytochromes as unique in sequence, they may not functionally be so. As shown in cytochrome nanowires, we may find that strict adherence to primary sequence is not as important for function as it is for other types of proteins. Based on this evidence, multi-heme cytochrome classification may need to be guided by a combination of local gene synteny and structural comparisons being made possible by the emergence of Cryo-EM and AlphaFold-like tools. We may be able to shrink this growing universe of sequence variation by clustering according to common shapes, heme packing motifs, and local organization patterns (45).

In conclusion, this study suggests caution when using annotation tools, and raises new questions about the perplexing “redundancy” of multi-heme cytochromes in *Shewanella* and *Geobacter*. Are increased mutation rates a result of a constant searching for more fit cytochromes, or are protein sequences simply drifting as they are not penalized as strongly as they are in enzymes? How much mutation can a multi-heme cytochrome tolerate before its function changes? Are enhanced mutation rates unique to specific extracellular electron transfer-performing organisms or are they a broader evolutionary trend in multi-heme cytochromes as a class? What is the size and mechanism supporting this this vast horizonal gene transfer pool? Answers to these questions will help address the severe limitations currently facing annotation of cytochrome-rich genomes, and help in identification of new organisms capable of extracellular electron transfer.

## Methods

### Selection and preparation of genomes for analysis

Genomes were obtained from NCBI in October 2021. We selected assemblies identified as ‘*Geobacter’* having at least 50X sequencing coverage and, if the genome was a metagenomic MAG, ≥ 3 MB to filter out low quality or incomplete assemblies. For each *Shewanella* species, a strain with a complete genome was selected. *Geobacter* and *Shewanella* genomes having no multi-heme cytochromes were removed from further analysis. The *Geobacter sulfurreducens* PCA genome was manually annotated to reflect the current literature on cytochrome and cytochrome related genes and then used as a reference to annotate our selected *Geobacter* genomes using Prokka (*19*). For *Shewanella* species *Shewanella oneidensis* MR-1 was used as the Prokka reference. In addition, all genes encoding mutiheme cytochromes were annotated with the number of hemes in case any lacked homologs in the reference strain, and to create the full set for pangenome analysis.

### Phylogenetic trees

Maximum likelihood trees for genomes were constructed with PhyloPhlAn using the amino acid supermatrix configuration file, which uses the PROTGAMMA-LG model of amino acid substitution, and the PhyloPhlAn database of 400 universal marker genes (*28*). For the *Geobacter* tree, the --medium diversity setting was used, as the selected genomes are now known to be from several genera in the family Geobacteraceae. *Desulfuromonas acetoxidans* DSM 684 and *Pelobacter carbinolicus* DSM 2380, from the Desulfuromonadaceae family, were used as the outgroup. High diversity was used for the combined *Geobacter* (Desulfobacterota) and *Shewanella* (Pseudomonadota) tree. *Geothrix fermentans* DSM 14018 and *Acidobacterium capsulatum* ATCC 51196 from Phylum Acidobacteriota were used as the outgroup.

For alignment of gene sequences the online MAFFT tool from EMBL-EBI was used with default parameters. Maximum likelihood gene trees were constructed using RAxML 8.2.11 (pthread), using the GTRGAMMA or PROTGAMMA models, depending on whether DNA or protein was being analyzed (*32*, *33*). RAxML was also used for bootstrapping, using rapid bootstrap analysis for protein-based genome trees and the PROTGAMMA-LG model. Both the *Geobacter* tree (Fig. 1) and the combined *Geobacter* and *Shewanella* tree (Figs. S3, S4) converged after 50 replicates. Rapid bootstrap analysis for the *omcSTJ* tree was performed with convergence after 600 replicates.

### Homolog clustering

Homolog clustering was performed with the bioinformatic tool get_homologues (*20*, *21*, *34–41*). Unless otherwise specified, genomes were analyzed using a reference genome and the OMCL algorithm, with all other settings as default. Pangenome matrices were obtained using the OMCL algorithm.

### Cytochrome diversity by phylogenetic distance

To calculate the number of new cytochromes identified with increasing phylogenetic distance from a reference genome, both get_homologues and a maximum likelihood *Geobacter* and *Shewanella* tree were used. Distances between each genome and their respective reference were calculated using the *Geobacter* and *Shewanella* maximum likelihood tree. Then get_homologues was used to analyze genomes in order of distance using inflation values (clustering stringency) of 1 (minimum stringency), 1.5 (default), and 5 (maximum stringency). The number of new cytochrome clusters per genome added was then calculated and plotted against phylogenetic distance.

### HMM construction and FeGenie

Input sequences for HMMs were chosen by get_homologues clustering or synteny. Amino acid sequences were aligned using MUSCLE (*42*). HMMs were constructed using hmmbuild in the bioinformatic package HMMER (*43*). Alignments and HMMs were manually added to FeGenie, along with our manually selected bitscore cutoff, to identify sequence hits (*22*).

### Mutation rate analysis

Input sequences were obtained from get_homologs cytochrome and “core” gene and protein homolog clusters. To ensure that comparisons were between homologs, cytochrome clusters were curated as follows: 1) the number of heme binding motifs must equal the reference ± 2, 2) the protein sequence length must be within 20% of the reference, 3) multiple sequences from the same genome within a cluster must be verified to be a duplication or the correct homolog identified, or else the sequences will be removed from further analysis. Protein sequences from homolog clusters were aligned using Clustal Omega (*26*). Sequences in gene homolog clusters were then codon formatted with their corresponding amino acid alignment files. Files with codon-formatted sequences were then converted to axt format using a custom script. Mutation rates for each homolog cluster were obtained by pairwise comparison of each sequence with the cluster’s PCA or MR-1 sequence using the modified Yang-Nielsen (MYN) method with the KaKs Calculator bioinformatic tool (*25*, *44*).

### Statistical analyses

All statistical analyses were performed in R (*45*). For all groups of dN/dS data, normality was tested using the Shapiro-Wilks test. Variance for normal data was calculated with the F-test. Cytochrome dN/dS values often had a non-normal distribution, with core dN/dS values being normal. In the former case these data were log transformed before performing two sample t-tests, or the Welch’s t-test, in the case of unequal variance. Where both sets of dN/dS values had non-normal distributions, the Wilcoxon Rank Sum test was used. For comparison of cytochrome dN/dS values by localization, the Kruskal-Wallis test was used with Dunn’s post-hoc test.

## Acknowledgements

This work was supported by a grant from the Office of Naval Research (N00014-18-1-2632) to JAG and DRB, and a grant from the Department of Energy to DRB (DE-SC0020322). The authors would like to thank Yaniv Brandvain for assistance with R.

## Supplemental Table & Figures

**Table S1.** Genomes used in bioinformatic analyses. All genomes and genome data were obtained from NCBI.

| Genomes | Genbank Accession Num | Isolation source |
| --- | --- | --- |
| *Geobacter anodireducens* SD-1 | GCA_001628815.1 | microbial fuel cell biofilm |
| *Geobacter argillaceus* ATCC_BAA_11 | GCA_007830735.1 | kaolin clays, USA |
| *Geobacter bemidjensis* Bem | GCA_000020725.1 | freshwater sediment, USA |
| *Geobacter bremsensis* R4 | GCA_014218275.1 | unknown |
| *Geobacter chapellei* DSM_13688 | GCA_018531195.1 | freshwater sediment, USA |
| *Geobacter daltonii* FRC-32 | GCA_000022265.1 | hydrocarbon contaminated sediment, USA |
| *Geobacter grbiciae* DSM_13689 | GCA_018531165.1 | freshwater sediment, USA |
| *Geobacter hydrogenophilus* DSM_136 | GCA_018501225.1 | hydrocarbon contaminated aquifer, USA |
| *Geobacter lovleyi* SZ | GCA_000020385.1 | freshwater sediment, South Korea |
| *Geobacter luticola* JCM_17780 | GCA_018476905.1 | lotus field sediment, Japan |
| *Geobacter metallireducens* GS-15 | GCA_000012925.1 | freshwater sediment, USA |
| *Geobacter pelophilus* Drf2 | GCA_002117975.1 | freshwater sediment, Germany |
| *Geobacter pickeringii* G13 | GCA_000817955.1 | kaolin clays, USA |
| *Geobacter soli* GSS01 | GCA_000816575.1 | humic layer soil, China |
| *Geobacter* sp. Bin_37_3 | GCA_009885845.1 | freshwater sediment, Canada |
| *Geobacter* sp. DSM_2909 | QAXR01000001.1 | starch wastewater digester, France |
| *Geobacter* sp. DSM_9736 | GCA_900187405.1 | paddy soil, Italy |
| *Geobacter* sp. FeAM09 | GCA_008330225.1 | forest soil, USA |
| *Geobacter* sp. GAC1 | GCA_003314535.1 | digester fed methanogenic reactor, USA |
| *Geobacter* sp. H2geo | GCA_003574885.1 | anode biofilm, USA |
| *Geobacter* sp. H5geo | GCA_009684525.1 | anode biofilm, USA |
| *Geobacter* sp. H7geo | GCA_009684515.1 | anode biofilm, USA |
| *Geobacter* sp. Jerry-YX | GCA_017338855.1 | petroleum soil, China |
| *Geobacter* sp. L1geo | GCA_003574895.1 | anode biofilm, USA |
| *Geobacter* sp. SVR | GCA_016865365.1 | mine soil, Japan |
| *Geobacter* sp. UBA698 | GCA_002298925.1 | anode biofilm |
| *Geobacter* sp. UBA2189 | GCA_002328035.1 | tailing pond, Canada |
| *Geobacter* sp. UBA6151 | GCA_002423105.1 | wastewater, Canada |
| *Geobacter* sp. UBA9964 | GCA_003490025.1 | groundwater |
| *Geobacter* sp. UBA12467 | GCA_003488785.1 | groundwater |
| *Geobacter sulfurreducens* PCA | GCA_000007985.2 | hydrocarbon contaminated sediment, USA |
| *Geobacter thiogenes* ATCC_BAA-34 | GCA_900167465.1 | contaminated subsoil, USA |
| *Geobacter uranireducens* Rf4 | GCA_000016745.1 | contaminated subsoil, USA |
| *Shewanella aestuarii* PN3F2 | GCF_011765625 | *Perinereis lineis* intestine |
| *Shewanella algae* RQs-106 | GCF_009730655 | activated sludge |
| *Shewanella amazonensis* SB2B | GCF_000015245 | marine sediment, Amazon River delta |
| *Shewanella avicenniae* FJAT-51800 | GCF_017354945 | estuary sediment, China |
| *Shewanella baltica* OS678 | GCF_000178875 | free-living marine, Baltic Sea |
| *Shewanella benthica* DB21MT-2 | GCA_900476435 | marine sediment, Mariana Trench |
| *Shewanella bicestii* JAB-1 | GCF_002216875 | bile, *Homo sapiens* |
| *Shewanella carassii* TUM17387 | GCF_019670705 | *Homo sapiens* |
| *Shewanella chilikensis* DC57 | GCA_011106835 | corroded pipe, Australia |
| *Shewanella cyperi* FJAT-53726 | GCF_017354925 | *Cyperus malaccensis* rhizosphere, China |
| *Shewanella decolorationis* Ni1-3 | GCA_007923045 | electroplating wastewater sludge |
| *Shewanella denitrificans* OS217 | GCF_000013765 | Baltic sea |
| *Shewanella dokdonensis* DSM_23626 | GCF_018394335 | seawater, South Korea |
| *Shewanella donghaensis* LT17 | GCF_007567505 | deep sea sediment, Sea of Japan |
| *Shewanella eurypsychrophilus* YLB-06 | GCF_007004545 | deep sea sediment, Indian Ocean |
| *Shewanella frigidimarina* NCIMB_400 | GCF_000014705 | seawater, North Sea |
| *Shewanella glacialis* TZS-4 | GCF_020511155 | sea ice, Baltic Sea |
| *Shewanella halifaxensis* HAW-EB4 | GCF_000019185 | marine sediment, Atlantic Ocean |
| *Shewanella inventiois* D1489 | GCF_019931735 | marine sediment, Pacific Ocean |
| *Shewanella japonica* KCTC_22435 | GCF_002075795 | seawater, Troitza Bay |
| *Shewanella khirikhana* TH2012 | GCF_003957745 | *Penaeus vannamei* hepatopancreas |
| *Shewanella litorisediminis* SMK1-12 | GCF_016834455 | tidal flat sediment, South Korea |
| *Shewanella livingstonensis* LMG_1986 | GCF_003855395 | Antarctic water |
| *Shewanella loihica* PV-4 | GCF_000016065 | hydrothermal vent, Pacific Ocean |
| *Shewanella marisflavi* EP1 | GCF_002215585 | marine sediment, China |
| *Shewanella maritima* D4-2 | GCF_004295345 | seawater, South Korea |
| *Shewanella oneidensis* MR-1 | GCA_000146165 | freshwater sediment, Lake Oneida |
| *Shewanella pealeana* ATCC_700345 | GCF_000018285 | *Loligo pealei* nidamental gland |
| *Shewanella piezotolerans* WP3 | GCF_000014885 | deep sea sediment, Pacific Ocean |
| *Shewanella polaris* SM1901 | GCF_006385555 | brown alga, Arctic Ocean |
| *Shewanella psychrophila* WP2 | GCF_002005305 | deep sea sediment, Pacific Ocean |
| *Shewanella psychropiezotolerans* YLB | GCF_007197555 | deep sea sediment, Indian Ocean |
| *Shewanella putrefaciens* ATCC_8071 | GCF_016406325 | butter |
| *Shewanella sedimentimangrovi* FJAT-1 | GCF_017354965 | estuary sediment, China |
| *Shewanella sediminis* HAW-EB3 | GCF_000018025 | marine sediment, Atlantic Ocean |
| *Shewanella vesiculosa* M7 | GCA_021560015 | marine sediment, Antarctic coast |
| *Shewanella violacea* DSS12 | GCF_000091325 | deep sea sediment, Ryukyu Trench |
| *Shewanella woodyi* ATCC_51908 | GCF_000019525 | detritus, Alboran Sea |
| *Shewanella xiamenensis* CQ-Y1 | GCA_019973655 | oilfield wastewater, China |

**Figure S1.**
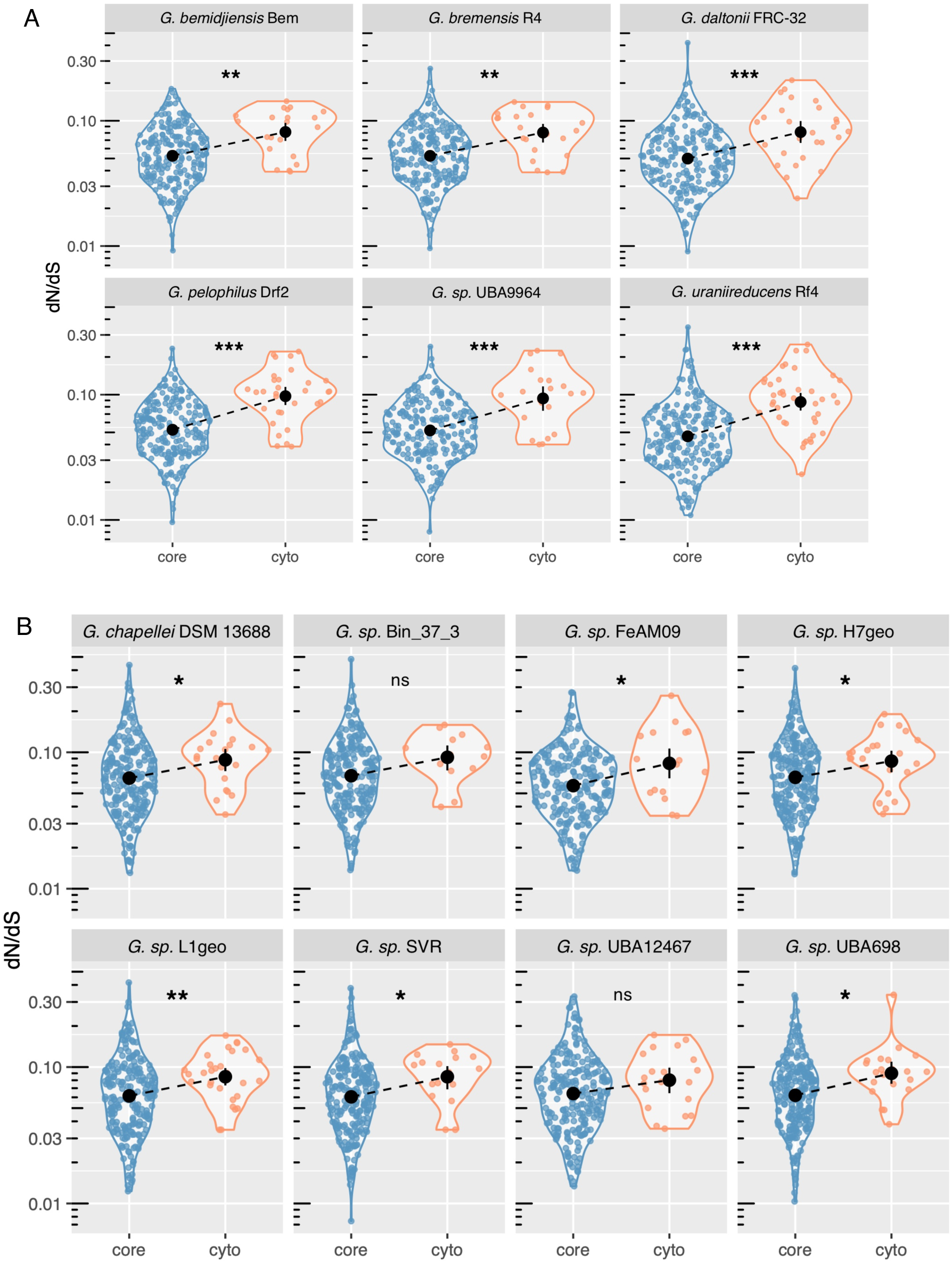
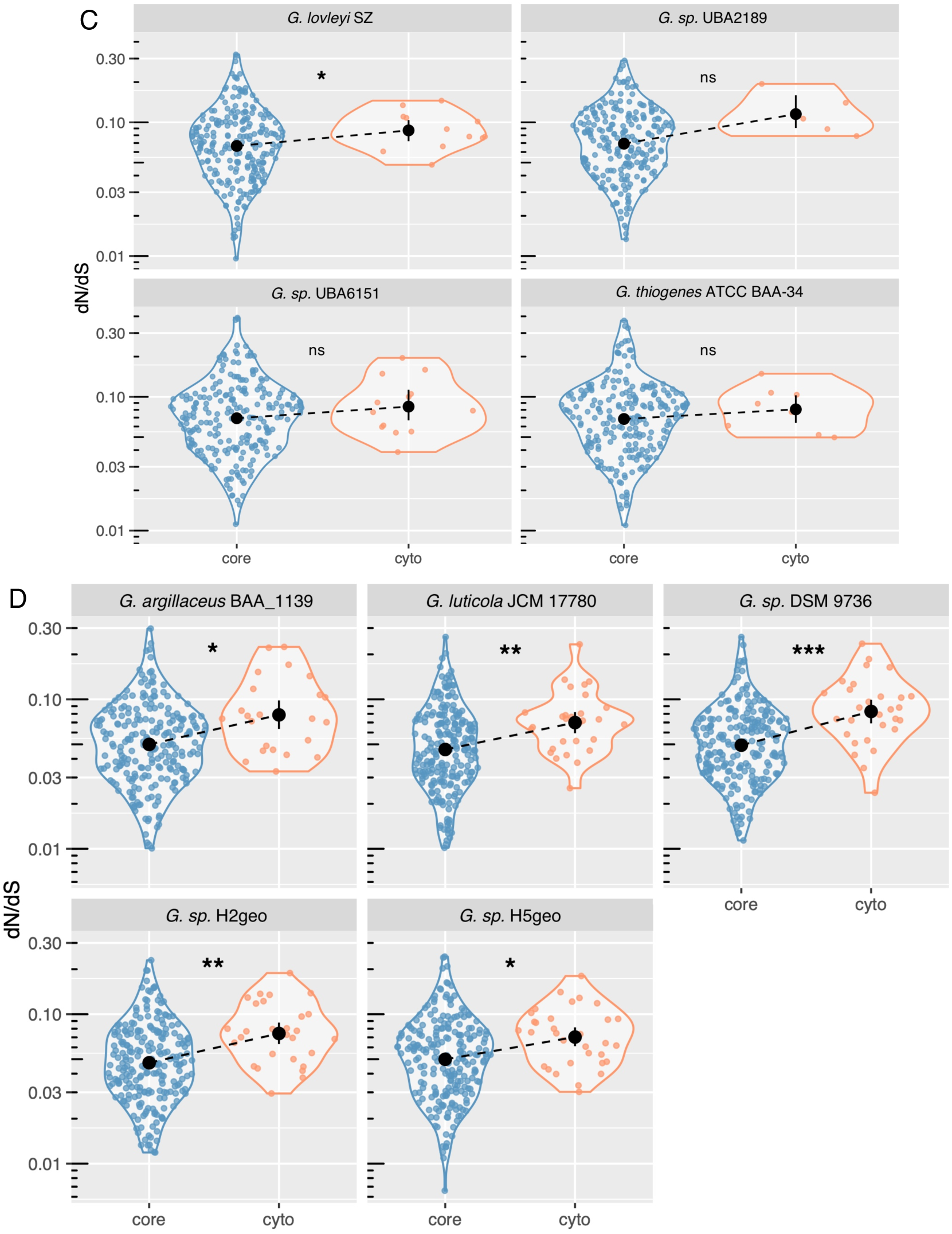
dN/dS analysis of core and MHC proteins in 23 *Geobacter* species, grouped phylogenetically. A) *G. bemidjiensis* Bem clade, B) *G. chapellei* DSM 13688 clade, C) *G. lovleyi* SZ clade, and D) *G. argillaceus* BAA_1139 clade. Significance: * p < 0.05, ** p < 0.01, *** p < 0.001, ns – not significant. *G. sp.* DSM 2909 and *G. sp.* GAC1 had too few MHCs and were omitted.

**Figure S2.**
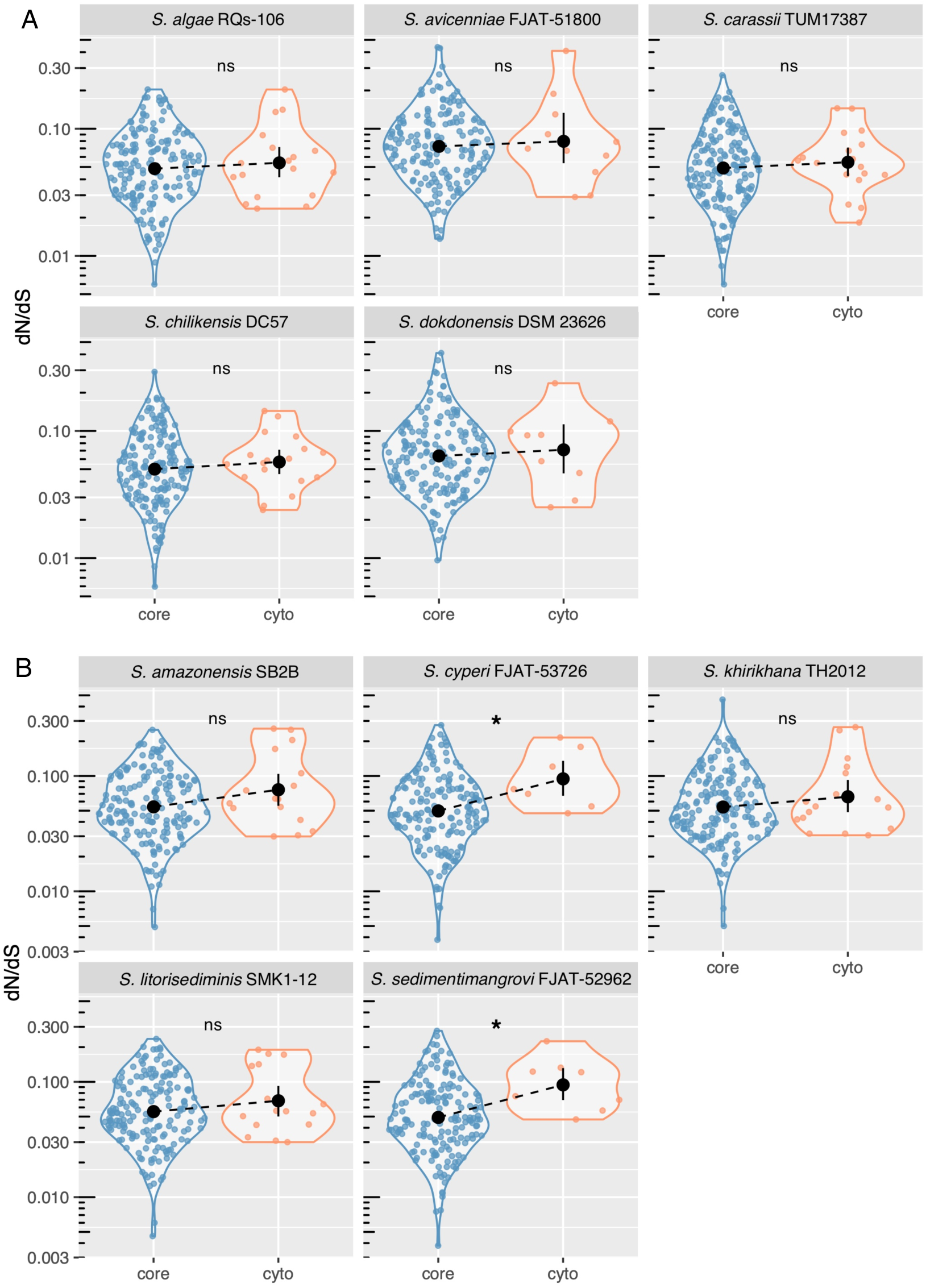
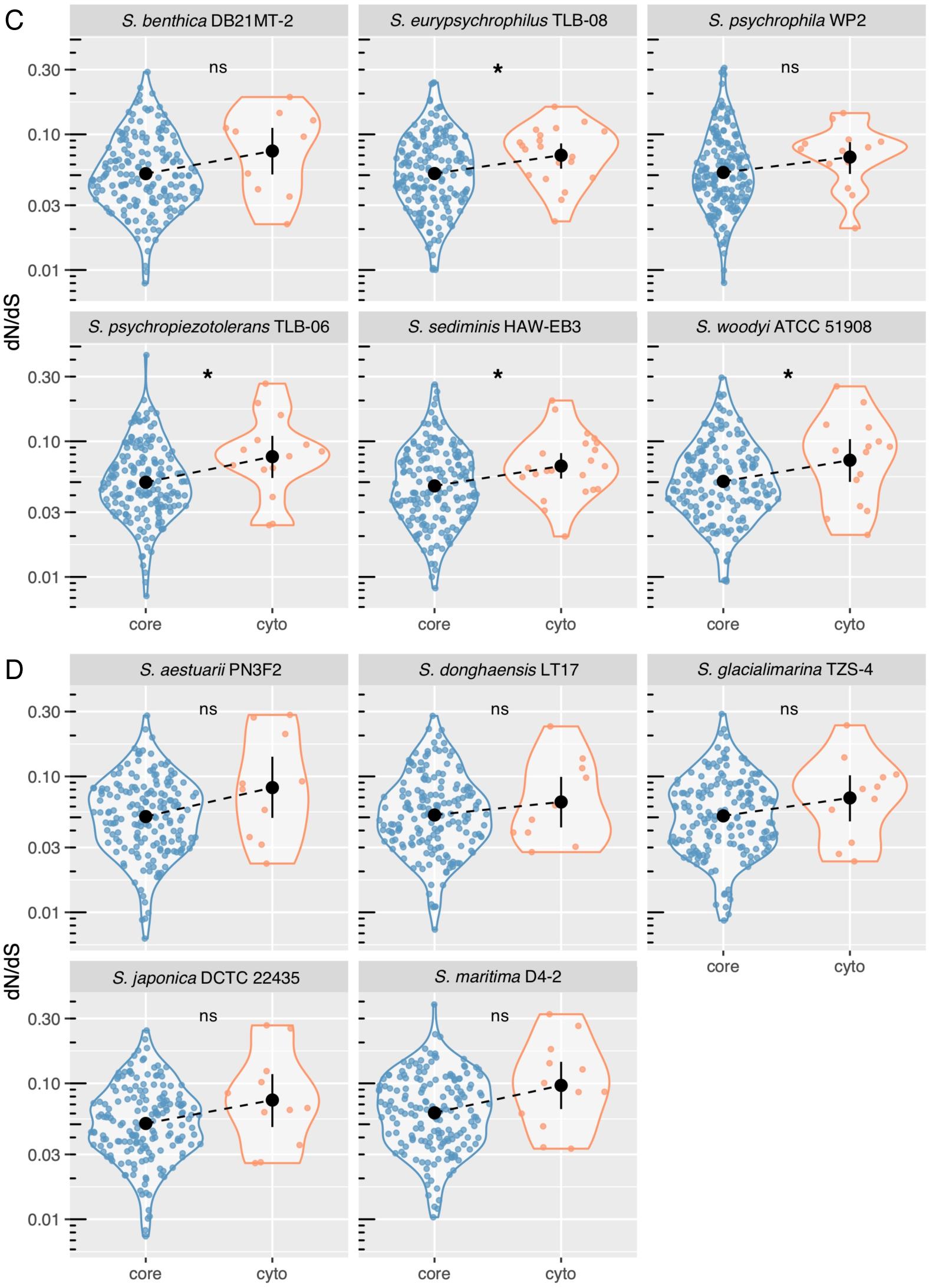
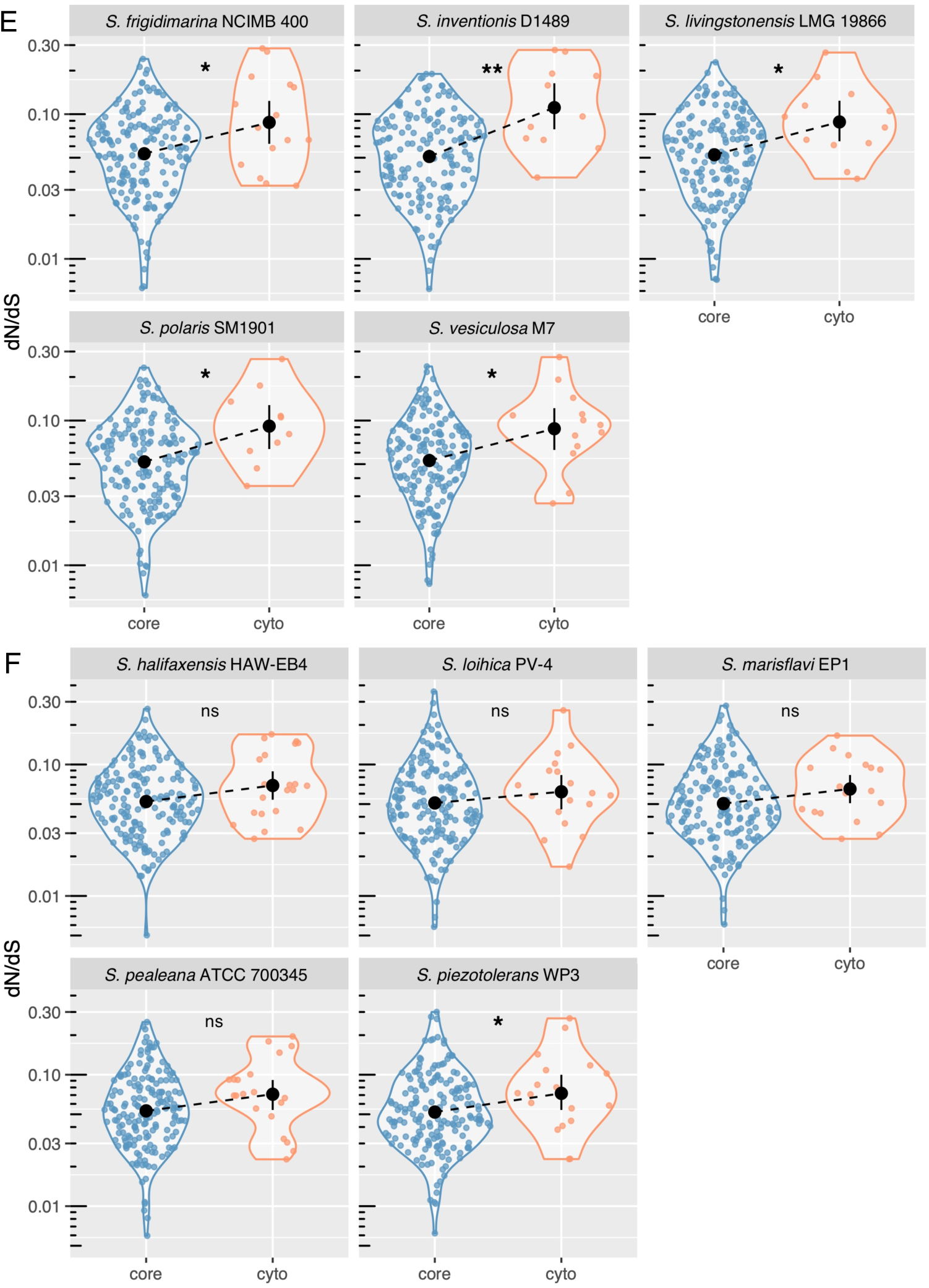
dN/dS analysis of core and MHC proteins in 31 *Shewanella* species, grouped phylogenetically. A) *S. algae* RQs-106 clade, B) *S. amazonensis* SB2B clade, C) *S. benthica* DB21MT-2 clade, D) *S. aestuarii* PN3F2 clade, E) *S. frigidimarina* NCIMB 400 clade, and F) *S. halifaxensis* HAW-EB4 clade. Significance: * p < 0.05, ** p < 0.01, ns – not significant

**Figure S3.**
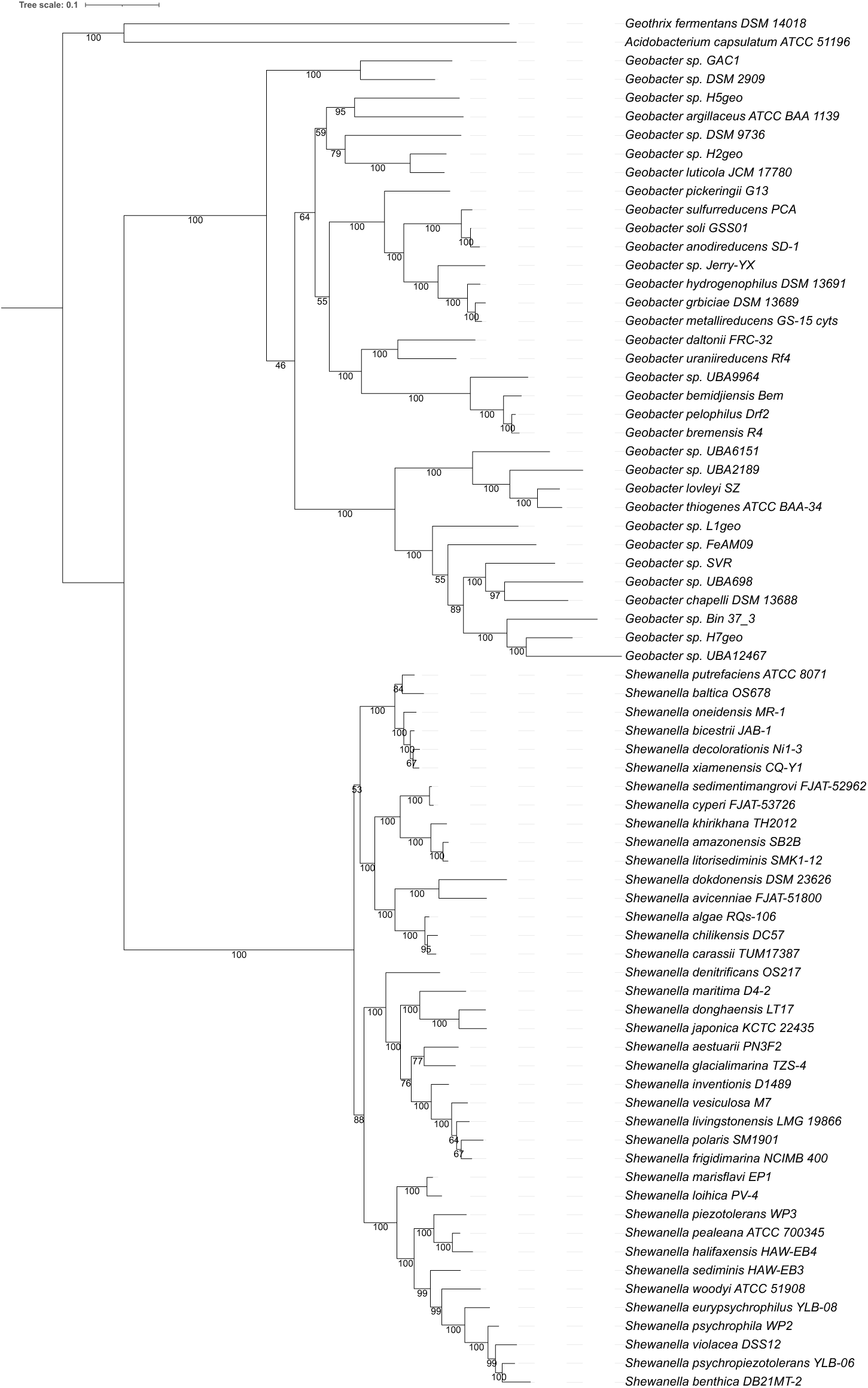
Combined Geobacter and Shewanella phylogenetic tree with bootstraps.

**Figure S4.**
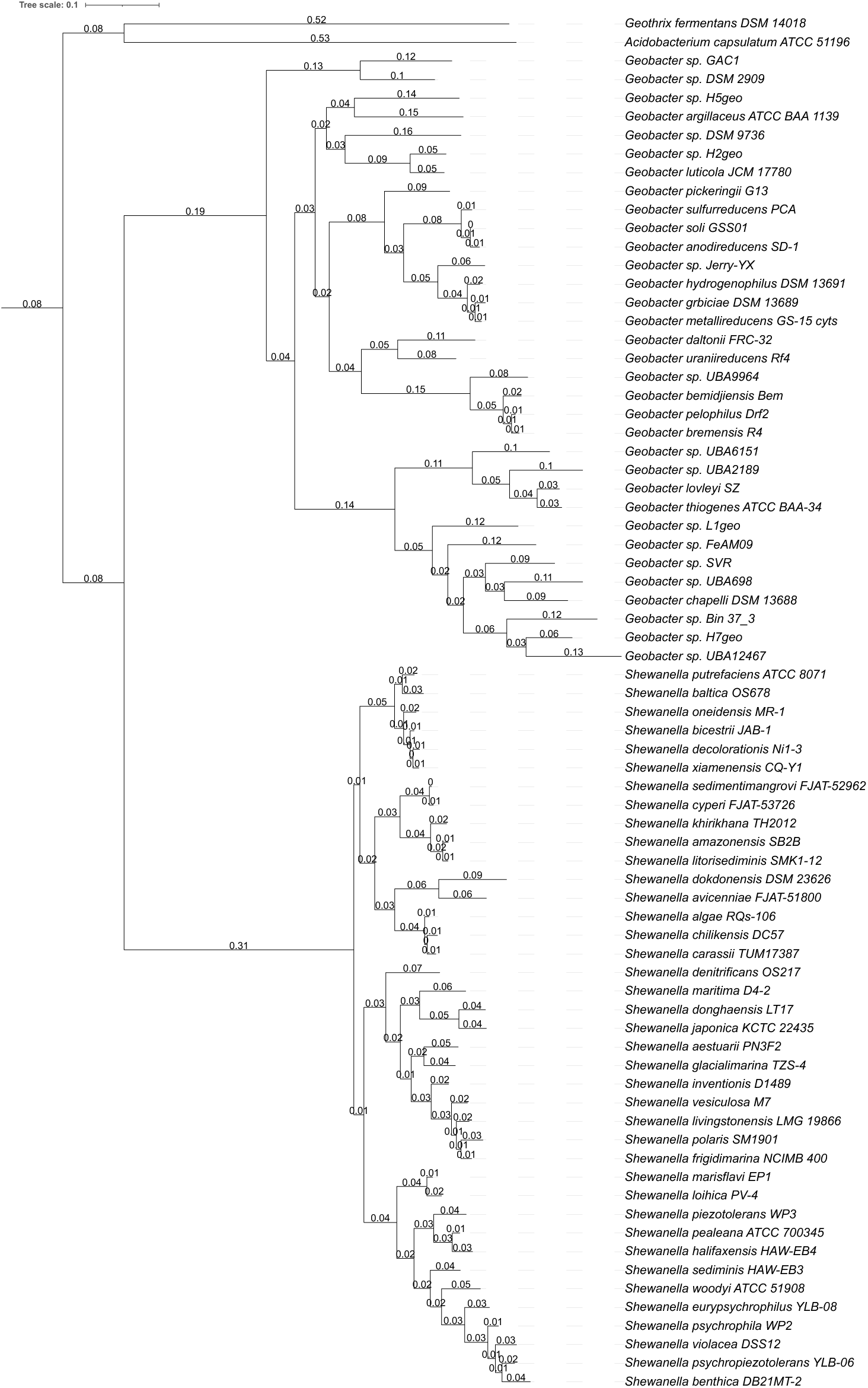
Combined Geobacter and Shewanella phylogenetic tree with branchlengths. Branchlength data was used to determine genome order and the x-axis scale in Fig. 9.

## References

1. C. M. Paquete, L. Morgado, C. A. Salgueiro, R. O. Louro, Molecular mechanisms of microbial extracellular electron transfer: the importance of multiheme cytochromes. Front. Biosci.-Landmark 27, 174 (2022).

2. U. Schröder, F. Harnisch, Life electric— Nature as a blueprint for the development of microbial electrochemical technologies. Joule 1, 244–252 (2017).

3. O. N. Lemaire, V. M. ’ Ejean, C. Iobbi-Nivol, The Shewanella genus: ubiquitous organisms sustaining and preserving aquatic ecosystems. FEMS Microbiol. Rev., 1–16 (2020).

4. G. L. Garbini, A. Barra Caracciolo, P. Grenni, Electroactive bacteria in natural ecosystems and their applications in microbial fuel cells for bioremediation: A review. Microorganisms 11, 1255 (2023).

5. D. R. Bond, Daniel R; Lovley, Electricity production by Geobacter sulfurreducens attached to electrodes. Appl. Environ. Microbiol. 69, 1548–1555 (2003).

6. A.-E. Rotaru, P. M. Shrestha, F. Liu, B. Markovaite, S. Chen, K. P. Nevin, D. R. Lovley, Direct interspecies electron transfer between Geobacter metallireducens and Methanosarcina barkeri. Appl. Environ. Microbiol. 80, 4599–605 (2014).

7. J. Zhou, J. A. Smith, M. Li, D. E. Holmes, Methane production by Methanothrix thermoacetophila via direct interspecies electron transfer with Geobacter metallireducens. mBio 0, e00360–23 (2023).

8. N. L. Ing, M. Y. El-Naggar, A. I. Hochbaum, Going the Distance: Long-Range Conductivity in Protein and Peptide Bioelectronic Materials. J. Phys. Chem. B 122, 10403–10423 (2018).

9. J. T. Atkinson, M. S. Chavez, C. M. Niman, M. Y. El-Naggar, Living electronics: A catalogue of engineered living electronic components. Microb. Biotechnol. 16, 507–533 (2022).

10. S. Choi, C. H. Chan, D. R. Bond, Lack of specificity in Geobacter periplasmic electron transfer. J. Bacteriol. 204, e00322–22 (2022).

11. P. R. Pokkuluri, Y. Y. Londer, N. E. C. Duke, W. C. Long, M. Schiffer, Family of cytochrome c7-type proteins from Geobacter sulfurreducens: structure of one cytochrome c7 at 1.45 Å resolution. Biochemistry 43, 849–859 (2004).

12. L. Morgado, M. Bruix, M. Pessanha, Y. Y. Londer, C. A. Salgueiro, Thermodynamic characterization of a triheme cytochrome family from Geobacter sulfurreducens reveals mechanistic and functional diversity. Biophys. J. 99, 293–301 (2010).

13. E. Howley, R. Krajmalnik-Brown, C. I. Torres, Cytochrome gene expression shifts in Geobacter sulfurreducens to maximize energy conservation in response to changes in redox conditions. Biosens. Bioelectron. 237, 115524 (2023).

14. C. E. Levar, C. H. Chan, M. G. Mehta-Kolte, D. R. Bond, An inner membrane cytochrome required only for reduction of high redox potential extracellular electron acceptors. mBio 5 (2014).

15. F. Wang, Y. Gu, J. P. O’Brien, S. M. Yi, S. E. Yalcin, V. Srikanth, C. Shen, D. Vu, N. L. Ing, A. I. Hochbaum, E. H. Egelman, N. S. Malvankar, Structure of microbial nanowires reveals stacked hemes that transport electrons over micrometers. Cell 177, 361–369.e10 (2019).

16. T. Mehta, M. V. Coppi, S. E. Childers, D. R. Lovley, Outer membrane c-type cytochromes required for Fe(III) and Mn(IV) oxide reduction in Geobacter sulfurreducens. Appl. Environ. Microbiol. 71, 8634–8641 (2005).

17. Z. Zhou, P. Q. Tran, A. M. Breister, Y. Liu, K. Kieft, E. S. Cowley, U. Karaoz, K. Anantharaman, METABOLIC: high-throughput profiling of microbial genomes for functional traits, metabolism, biogeochemistry, and community-scale functional networks. Microbiome 10, 33 (2022).

18. J. R. Lloyd, C. Leang, A. L. Hodges Myerson, M. V. Coppi, S. Cuifo, B. Methe, S. J. Sandler, D. R. Lovley, Biochemical and genetic characterization of PpcA, a periplasmic c-type cytochrome in Geobacter sulfurreducens. Biochem. J. 369, 153–161 (2003).

19. T. Seemann, Prokka: rapid prokaryotic genome annotation. Bioinformatics 30, 2068–2069 (2014).

20. B. Contreras-Moreira, P. Vinuesa, GET_HOMOLOGUES, a versatile software package for scalable and robust microbial pangenome analysis. Appl. Environ. Microbiol. 79, 7696– 7701 (2013).

21. L. Li, C. J. Stoeckert, D. S. Roos, OrthoMCL: Identification of ortholog groups for eukaryotic genomes. Genome Res. 13, 2178–2189 (2003).

22. A. I. Garber, K. H. Nealson, A. Okamoto, S. M. McAllister, C. S. Chan, R. A. Barco, N. Merino, FeGenie: a comprehensive tool for the identification of iron genes and iron gene neighborhoods in genome and metagenome assemblies. Front. Microbiol. 11 (2020).

23. M. J. Edwards, D. J. Richardson, C. M. Paquete, T. A. Clarke, Role of multiheme cytochromes involved in extracellular anaerobic respiration in bacteria. Protein Sci. Publ. Protein Soc. 29, 830–842 (2020).

24. L. D. Hurst, The Ka/Ks ratio: diagnosing the form of sequence evolution. Trends Genet. 18, 486–487 (2002).

25. Z. Zhang, KaKs_Calculator 3.0: Calculating selective pressure on coding and non-coding sequences. Genomics Proteomics Bioinformatics 20, 536–540 (2022).

26. F. Sievers, A. Wilm, D. Dineen, T. J. Gibson, K. Karplus, W. Li, R. Lopez, H. McWilliam, M. Remmert, J. Söding, J. D. Thompson, D. G. Higgins, Fast, scalable generation of high-quality protein multiple sequence alignments using Clustal Omega. Mol. Syst. Biol. 7, 539 (2011).

27. B. Schuetz, M. Schicklberger, J. Kuermann, A. M. Spormann, J. Gescher, Periplasmic electron transfer via the c-type cytochromes MtrA and FccA of Shewanella oneidensis MR-1. Appl. Environ. Microbiol. 75, 7789–7796 (2009).

28. F. Asnicar, A. M. Thomas, F. Beghini, C. Mengoni, S. Manara, P. Manghi, Q. Zhu, M. Bolzan, F. Cumbo, U. May, J. G. Sanders, M. Zolfo, E. Kopylova, E. Pasolli, R. Knight, S. Mirarab, C. Huttenhower, N. Segata, Precise phylogenetic analysis of microbial isolates and genomes from metagenomes using PhyloPhlAn 3.0. Nat. Commun. 11, 2500 (2020).

29. J. E. Butler, N. D. Young, M. Aklujkar, D. R. Lovley, Comparative genomic analysis of Geobacter sulfurreducens KN400, a strain with enhanced capacity for extracellular electron transfer and electricity production. BMC Genomics 13, 471–471 (2012).

30. F. Wang, K. Mustafa, V. Suciu, K. Joshi, C. H. Chan, S. Choi, Z. Su, D. Si, A. I. Hochbaum, E. H. Egelman, D. R. Bond, Cryo-EM structure of an extracellular Geobacter OmcE cytochrome filament reveals tetrahaem packing. Nat. Microbiol., 1–10 (2022).

31. A. W. Grenfell, P. J. Intile, J. A. McFarlane, D. C. Leung, K. Abdalla, M. C. Wold, E. D. Kees, J. A. Gralnick, The outer membrane cytochrome OmcA Is essential for infection of Shewanella oneidensis by a zebrafish-associated bacteriophage. J. Bacteriol. 205, e00469–22 (2023).

32. K. Katoh, K. Misawa, K. Kuma, T. Miyata, MAFFT: a novel method for rapid multiple sequence alignment based on fast Fourier transform. Nucleic Acids Res. 30, 3059–3066 (2002).

33. A. Stamatakis, RAxML version 8: a tool for phylogenetic analysis and post-analysis of large phylogenies. Bioinformatics 30, 1312–1313 (2014).

34. P. Vinuesa, B. Contreras-Moreira, “Robust identification of orthologues and paralogues for microbial pan-genomics using GET_HOMOLOGUES: a case study of pIncA/C plasmids” in Bacterial Pangenomics: Methods and Protocols, A. Mengoni, M. Galardini, M. Fondi, Eds. (Springer, New York, NY, 2015; 10.1007/978-1-4939-1720-4_14)Methods in Molecular Biology, pp. 203–232.

35. B. Buchfink, C. Xie, D. H. Huson, Fast and sensitive protein alignment using DIAMOND. Nat. Methods 12, 59–60 (2015).

36. J. E. Stajich, D. Block, K. Boulez, S. E. Brenner, S. A. Chervitz, C. Dagdigian, G. Fuellen, J. G. R. Gilbert, I. Korf, H. Lapp, H. Lehväslaiho, C. Matsalla, C. J. Mungall, B. I. Osborne, M. R. Pocock, P. Schattner, M. Senger, L. D. Stein, E. Stupka, M. D. Wilkinson, E. Birney, The Bioperl toolkit: Perl modules for the life sciences. Genome Res. 12, 1611–1618 (2002).

37. S. F. Altschul, T. L. Madden, A. A. Schäffer, J. Zhang, Z. Zhang, W. Miller, D. J. Lipman, Gapped BLAST and PSI-BLAST: a new generation of protein database search programs. Nucleic Acids Res. 25, 3389–3402 (1997).

38. R. D. Finn, P. Coggill, R. Y. Eberhardt, S. R. Eddy, J. Mistry, A. L. Mitchell, S. C. Potter, M. Punta, M. Qureshi, A. Sangrador-Vegas, G. A. Salazar, J. Tate, A. Bateman, The Pfam protein families database: towards a more sustainable future. Nucleic Acids Res. 44, D279–D285 (2016).

39. N. P. Brown, C. Leroy, C. Sander, MView: a web-compatible database search or multiple alignment viewer. Bioinformatics 14, 380–381 (1998).

40. hmmscan :: search sequence(s) against a profile database HMMER 3.1b2, (2015); http://hmmer.org.

41. D. M. Kristensen, L. Kannan, M. K. Coleman, Y. I. Wolf, A. Sorokin, E. V. Koonin, A. Mushegian, A low-polynomial algorithm for assembling clusters of orthologous groups from intergenomic symmetric best matches. Bioinformatics 26, 1481–1487 (2010).

42. R. C. Edgar, MUSCLE: multiple sequence alignment with high accuracy and high throughput. Nucleic Acids Res. 32, 1792–1797 (2004).

43. S. R. Eddy, HMMER, version 3.3.2; www.hmmer.org.

44. Z. Zhang, J. Li, J. Yu, Computing Ka and Ks with a consideration of unequal transitional substitutions. BMC Evol. Biol. 6, 44 (2006).

45. R Core Team, R: A language and environment for statistical computing., version RStudio 2021.09.0, R Foundation for Statistical Computing (2021); www.R-project.org.

45. Soares, R., Fonseca, B.M., Nash, B.W. et al. A survey of the *Desulfuromonadia* “cytochromome” provides a glimpse of the unexplored diversity of multiheme cytochromes in nature. BMC Genomics 25, 982 (2024).

46. Shen, C., Salazar-Morales, A.I., Jung, W., Erwin, J., Gu, Y., Coelho, A., Gupta, K., Yalcin, S.E., Samatey, F.A., Malvankar, N.S. A widespread and ancient bacterial machinery assembles cytochrome OmcS nanowires essential for extracellular electron transfer. Cell Chemical Biology (2025).

